# A consensus spinal cord cell type atlas across mouse, macaque, and human

**DOI:** 10.64898/2026.02.04.703852

**Authors:** Matthew T. Schmitz, Nelson J. Johansen, Niklas Kempynck, Inkar Kapen, Yuanyuan Fu, Madeleine Hewitt, Stephanie C. Seeman, Emily Kussick, Olivia Gautier, Michael J. Leone, Song-Lin Ding, Yuan Gao, Ashwin Bhandiwad, Jeanelle Ariza, Angela Ayala, Stuard Barta, Jacob A. Blum, Liliana Cano-Gomez, Trangthanh Cardenas, Anish Bhaswanth Chakka, Nasmil Valera Cuevas, Nicholas Donadio, Katherine A. Fancher, Emma D. Thomas, Rebecca Ferrer, Jeff Goldy, Samantha D. Hastings, Daniel Hirschstein, Windy Ho, Cindy Huang, Zoe C. Juneau, Sa Rang Kim, Zachary R. Lewis, Elizabeth Liang, Naomi X Martin, Josh Nagra, Dakota Newman, Myung-Chul Noh, Paul Olsen, Alana Oyama, Nick Pena, Helen Poldsam, Patrick L. Ray, Melissa Reding, Christine Rimorin, Augustin Ruiz, Nadiya V. Shapovalova, Lyudmila Shulga, Cassandra Sobieski, Amy Torkelson, Morgan Wirthlin, Shenqin Yao, Helen Lai, Andreas Pfenning, Allan-Hermann Pool, Rebecca Seal, Bosiljka Tasic, Jonathan T. Ting, Jack Waters, Zizhen Yao, Aaron D. Gitler, Delissa McMillen, Lydia Ng, Hongkui Zeng, Cindy T. J. van Velthoven, Tanya L. Daigle, Kimberly A. Smith, Rebecca D. Hodge, Ed S. Lein, Trygve E. Bakken

## Abstract

The spinal cord contains evolutionarily conserved cell types critical for motor function, sensory processing, and autonomic regulation, many of which are implicated in diverse neurological diseases and injuries. Yet the field lacks a comprehensive molecular characterization of cellular diversity in human, macaque, and mouse spinal cord. Here, we present a unified, cross-species cell type atlas based on the integration of single-nucleus gene expression, chromatin accessibility, and spatial transcriptomic data from segments within cervical, thoracic, lumbar, and sacral regions, including motor neurons (MNs) sampled across the entire rostro-caudal axis of the macaque spinal cord. Leveraging the spatial distributions of our molecularly defined cell types, we generated a cell type-guided anatomical map of spinal cord laminae and nuclei. We identified both conserved and species-specific cellular features, including gene expression patterns across distinct MN subtypes in the primate spinal cord. Cross-species cis-regulatory analysis and deep learning sequence models dissected the enhancer logic underlying viral targeting, uncovering conserved transcription factor grammar encoding cellular identity. Together, these results establish a unifying molecular and anatomical taxonomy of spinal cord cell types across species.

## INTRODUCTION

The mammalian spinal cord serves as a central hub linking the brain to the body, integrating sensory input, motor output, respiratory control, and autonomic regulation. Through its complex organization of neurons and glia, the spinal cord generates complex encodings of sensory information that are routed to the brain, while also enabling both rapid reflexive responses and fine motor control, as well as visceral organ homeostasis. These physiological processes are mediated by diverse and highly specialized cell types with distinct molecular and morphological features, which give rise to precise functional circuits. Motor systems show clear differences between species, with many more monosynaptic cortico-spinal control of MNs in the primate than in the mouse ^1^. In addition, α-MNs within the ventral horn drive voluntary movement and are selectively vulnerable in human neurodegenerative disorders such as amyotrophic lateral sclerosis (ALS). Within and across species, neurons vary in morphology, connectivity and function along the length of the cord ^2–4^ while molecular studies have shown broad conservation of cell types in the human and mouse dorsal horn ^5,6^. This suggests that functional differences likely arise from more subtle differences in molecular profiles or relative abundances of cell types.

Over the past decade, single-cell transcriptomic and spatial technologies have transformed our understanding of nervous system organization. Large-scale brain atlases have provided a major step towards standardized, universal interpretations of cell types in individual species, including human ^7^ and mouse ^8^. Additionally, comparative studies of primate cortex have established robustly conserved taxonomies ^9–11^, demonstrating that widespread cell type conservation underlies core circuit functions with some lineage-specific molecular specializations. These studies highlight the value of integrating transcriptomic and anatomical information at scale to understand the organization and evolutionary trajectories of the central nervous system.

In contrast, mammalian spinal cord atlases remain comparatively sparse. Previous studies in mouse have provided good coverage of cell types or segments ^12–14^, and recent human lumbar cord profiling identified major neuronal and non-neuronal populations ^6^, although ventral cell type homologies with mouse were less resolved and few MNs were profiled ^15^. A comparative study that was restricted to the dorsal horn integrated spatial, transcriptomic, and epigenomic data from human, macaque, and mouse to identify conserved cell types with a subset linked to genetic risk for chronic pain ^5^. However, these studies lack deep neuronal sampling across species and segments along the rostral-caudal axis, and there remains a need for a comprehensive atlas of cellular diversity in human and non-human primate spinal cord and cell type homologies with mouse.

Comparative spinal cord atlasing should connect conserved cell identities to the regulatory sequence logic that specifies cell type-selective chromatin accessibility across species. Coupling cross-species cCRE maps with sequence-based inference can expose conserved motif grammar and prioritize candidate regulatory elements most likely to generalize to primate and human spinal cord cell populations. Here, we present a cross-species single-nucleus and spatial atlas of the spinal cord for human, pig-tailed macaque, and mouse, integrating transcriptomic and epigenomic modalities. We profiled spinal cords from multiple individuals for each species using the 10x Genomics Multiome workflow to jointly measure chromatin accessibility and gene expression. Furthermore, we applied MERFISH spatial transcriptomics to examine the distribution of cell types in four spinal segments in two macaque animals. Our dataset encompasses cervical and lumbar levels from human donors and representation from cervical, thoracic, lumbar and sacral regions from macaque and mouse, thereby providing a level of coverage not achieved in previous studies. Across species, this resource comprises 49,239 neurons in macaque, 3,491 neurons in human, and 53,862 neurons in mouse, providing a unified, multimodal foundation for comparative analysis of spinal cord cell types and regulatory programs.

## RESULTS

### Cross-species consensus atlas of the spinal cord

To construct a cross-species consensus atlas of transcriptional and chromatin accessibility states in the adult spinal cord, we generated single-nucleus multiomic profiles from human and pig-tailed macaque (*Macaca nemestrina*) and integrated these data with a mouse spinal cord data set described in a companion study (Gao et al. 2026). We collected post-mortem spinal cord tissue from six macaques and three human donors. From macaque, we sampled the four major rostrocaudal regions (cervical, thoracic, lumbar, and sacral), and in human we focused on cervical and lumbar segments. We enriched for neuronal (NeuN+) nuclei using fluorescence-activated nuclei sorting (FANS) and profiled paired transcriptome and chromatin accessibility data from 100,895 macaque nuclei and 21,388 human nuclei using 10x Genomics multiome. To further enrich for MNs we used FANS to capture BCL6+ neurons from the entire spinal cord of one macaque donor (Figure S1.2) ^16^. These datasets were integrated with 106,297 nuclei from a companion mouse multiome study spanning cervical through sacral spinal cord regions, generated using the same molecular workflow (Gao et al. 2026).

We generated a consensus taxonomy to enable comparisons of cellular abundances and molecular profiles across species. We applied scVI to integrate transcriptomic data from the three species and removed donor and species effects. Consensus clustering of these cross-species shared latent space grouped cells into 257 clusters (Figure S1). We used a recent human lumbar spinal cord taxonomy ^6^ to map prior labels onto our data providing an initial taxonomy. We organized the 257 transcriptomic clusters into a hierarchical cell type taxonomy based on conserved molecular, spatial, and developmental evidence. This unified hierarchy placed neurons and non-neuronal populations into 5 Classes representing neurotransmitter identity, 20 Subclasses representing neurotransmitter plus gross anatomical signature, and 49 Groups informed by published markers and nomenclature derived from Yadav et al. 2023 (Figure 1B-C, Figure S1.2).

**Figure 1.**
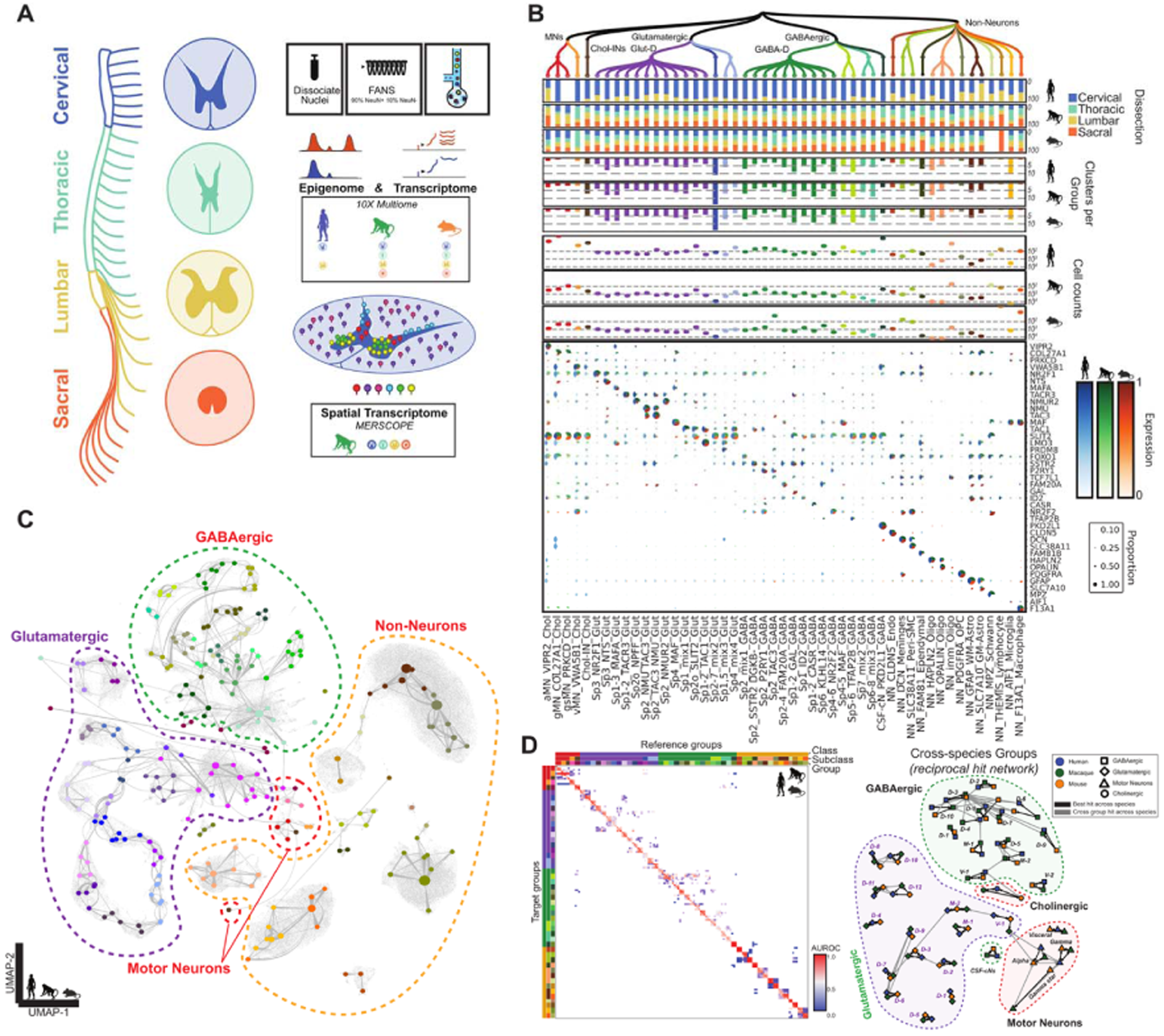
A consensus taxonomy of spinal cord cell types across human, macaque, and mouse. **(A)** Schematic of the spinal cord segments and genomics profile strategies across species. **(B)** Top, the transcriptomic taxonomy tree of Groups. Barplots represent the number of clusters within each Group for each species. Dotplots represent the number of cells supporting each Group from each species. Pie dot-plot of the species-specific expression for selected conserved marker genes per Group term, where color palettes represent expression in each species, and dot diameter represents the proportion of cells in each Group with non-zero counts for that gene. **(C)** Cross-species integrated constellation and UMAP representation of all consensus clusters colored by Group and linked based on expression similarity across species. **(D)** MetaNeighbor heatmap of ‘one vs best’ AUROC scores between Group terms across species using the conserved marker gene panel. MetaNeighbor network of ‘one vs best’ reciprocal hits (edges) between Groups (nodes) across species. Edges in which the reference group does not match the target group are colored by off-diagonal AUROC scores.

At the Group level, we found neuronal and non-neuronal homologous cell types or ‘orthotypes’ that were conserved between primates and mouse, and this was a similar level of conservation to cortical and subcortical brain regions ^9,17–22^ (Figures 1C-D and S1). To align the Group names with their molecular and spatial identity, we propose a tripartite nomenclature consisting of spatial distribution, marker genes, and a neurotransmitter or non-neuronal label that aligns with recent atlas work in mouse brain^8^ and primate basal ganglia ^17^. Markers were chosen that were conserved between primates and mouse in this study and also aligned with a companion mouse study (Cano Gomez et al. 2026) which was aligned to this taxonomy (Table S1).

Not surprisingly, as part of the central nervous system, the spinal cord contains non-neuronal Groups that are distributed across the brain. We identified 14 non-neuronal Groups from 9 Subclasses: Ependymal, Vascular, Astrocyte (Astro), Endothelial (Endo), Microglia, Macrophage, Oligodendrocyte Progenitor Cell (OPC), Oligodendrocyte (Oligo), and Schwann cells. Four of these groups (NN_MPZ_Schwann, NN_F13A1_Macrophage, NN_THEMIS_Lymphocyte, and NN_DCN_Meninges) are likely associated with peripheral nerves, blood vessels and meninges outside of the spinal cord. We identified Groups of astrocytes and oligodendrocytes which were more associated with white matter (NN_HAPLN_Oligo and NN_GFAP_WM-Astro) and with grey matter (NN_OPALIN_Oligo, NN_SLC7A10_GM-Astro). This is consistent with specializations of these non-neuronal populations in grey matter and white matter compartments of the basal ganglia^23^.

We identified 35 distinct Groups of neurons that were organized into 4 Classes (GABAergic, Glutamatergic, Cholinergic and Motor Neurons) and 10 Subclasses: dorsal, mid and ventral GABAergic neurons (GABA-D, GABA-M, GABA-V); dorsal, mid, ventral, and CSF-contacting Glutamatergic neurons (Glut-D, Glut-M, Glut-V, CSF-cNs); Cholinergic interneurons (Chol-IN); and skeletal and visceral Motor Neurons (Skeletal-MN, Visceral-MN). In general, GABA-D and Glut-D Groups were distinct, while the mid and ventral Groups contained significant underlying diversity. Some of these Groups (particularly ventral) were labeled “mix” because they include multiple cell types that could not be parsed, potentially due to technical noise resulting from dissociation of adult primate neurons or cross-species integration masking regional axes of variation. For example, Sp1,5_mix3_Glut contains clear subtypes that express prodynorphin (PDYN) which is associated with pain processing^5^. These Groups are described in more detail in a companion study of mouse spinal cord (Cano Gomez et al. 2026) (Table S1). We captured *bona fide* skeletal and visceral MNs from macaque, including skeletal subtypes alpha (aMN_VIPR2_Chol), gamma (gMN_COL27A14_Chol), and gamma* (gsMN_PRKCD_Chol) (Figures 1B and S2). These MNs expressed established markers including CHAT, ISL1/2 and BCL6 at higher levels, expressed more transcripts per cell than other neurons, and clustering divided the cells into groups clearly corresponding to skeletal and visceral motorneuron cell types (Figure 1, Table S2). By integrating our macaque data with external mouse datasets^12–14^, we further subdivided MN diversity into slow and fast firing alpha MNs (aMN-sf_PRKCD and aMN-ff_CHODL), gamma and gamma* MNs (gMN_DKK2 and gsMN_STXBP6), and 9 subtypes of visceral MNs, including rostral thoracic-specific GPC3+ vMNS^13^ and SST+ pelvic-ganglion innervating vMNs ^24^ (Figure S1.2). To our knowledge, this represents the first molecular profiling of these fine subdivisions of MNs in a primate.

### Anatomical distributions of cell types across the macaque spinal cord

We next defined a 300 gene MERFISH panel using markers from our snRNA-seq data and previously reported MN markers and examined the cytoarchitecture of the macaque spinal cord using spatial transcriptomics(Table S3) ^14^. We profiled 11 sections from 2 donors across 5 spinal cord segments (C3, C5, T4, L3, and S3)(Figure 2A) and annotated cells in the spatial data by mapping Classes, Subclasses, Groups and consensus clusters using MapMyCells (RRID:SCR_024672).

**Figure 2.**
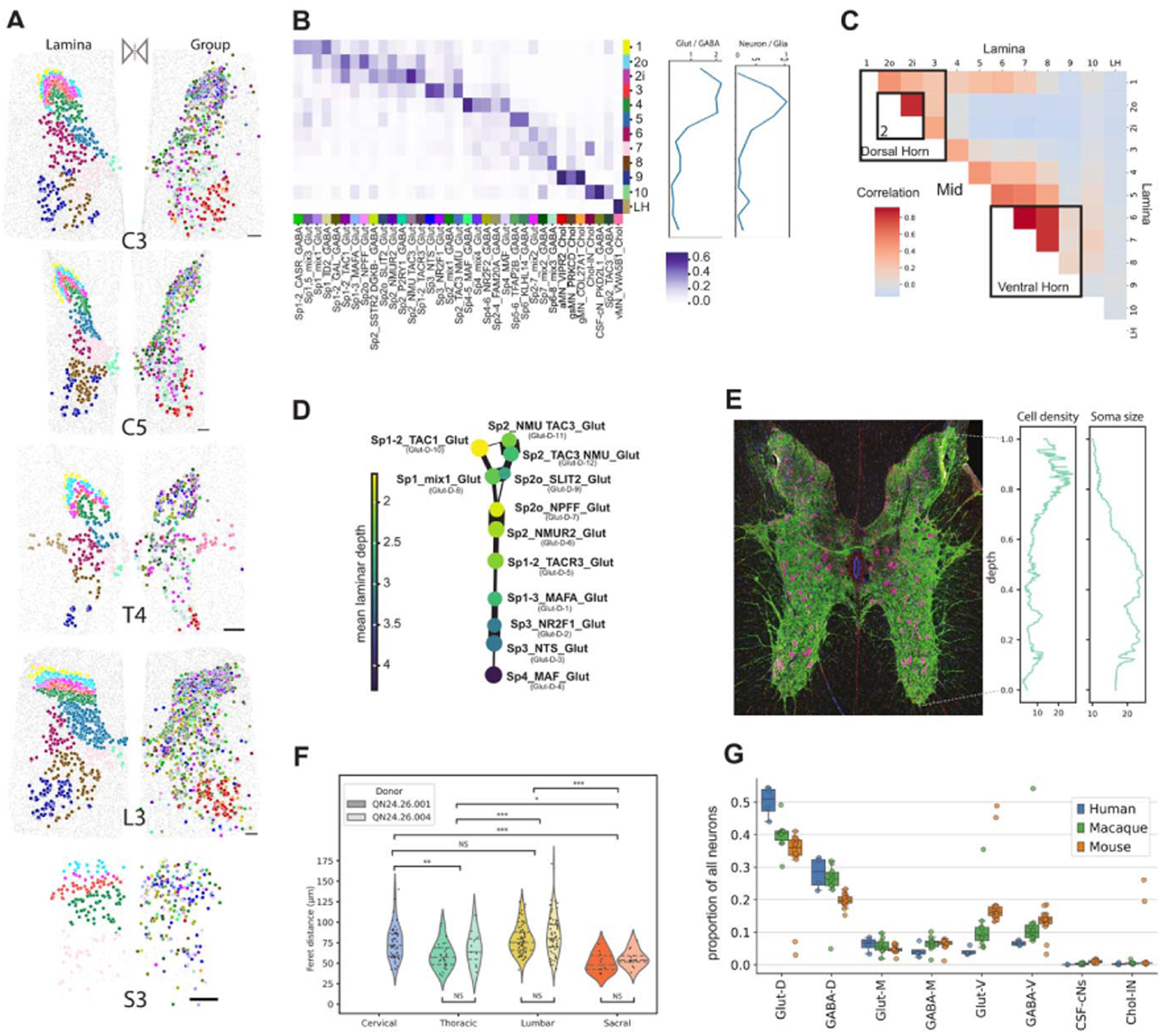
Anatomical distributions of cell types across the macaque spinal cord. **(A)** Macaque MERFISH data showing a representative single section’s hemisphere for each of the segments sampled, reflected to show manual annotations of approximate laminae (left) and Group taxonomic annotations (right). See **B** for Group color legend. Scale bars represent 300 um. **(B)** Approximate laminar distributions of neuron Groups, excluding Sacral sections. Values are cell counts, normalized by cells per layer, then normalized per group. Line plots on right show Glutamatergic / GABAergic neuron counts per lamina, and Neuron / Glia counts among each lamina’s neurons and their nearest neighbors. **(C)** Heatmap showing Pearson correlation of cell type composition vectors per lamina. **(D)** PAGA graph of Groups where edge thickness and point distance represents numbers of nearest neighbors in integrated scVI latent space, colored by the laminar depth (mean lamina value of cells in each Group, where Sp1 is 1, Sp2o is 1.66 and Sp2i is 2.33, Sp3 is 3, etc). **(E)** Immunofluorescence image of a thoracic section (tandem section to MERFISH from donor QN24.26.004) showing NeuN (magenta), DAPI (blue) and GFAP (green). Line plots on right show running mean of cell density (number of cells within 100 µm radius) and soma size (Feret diameter, µm). **(F)** Violin plots with overlaid stripplots showing Feret diameters of MNs from each donor in each segment **(G)** Boxplot with overlaid stripplot colored by species showing the proportion of each neuronal Subclass observed in each multiome data batch, highlighting species differences.

Using our mapped cell labels, we manually segmented regions of distinct neuron composition, which corresponded closely to Rexed laminae, except in sacral cord where laminae are less well defined. In cervical, thoracic and lumbar segments, all neuron Groups were enriched in laminae, with most localized to only one or two laminae, including well known Sp2_NMU TAC3_Glut and Sp3_NTS_Glut in lamina 2 and 3 respectively (Figure 2B). Based upon cell type composition, laminae Sp1-3 are similar and constitute the dorsal horn, while Sp4, Sp5, and Sp10 are each distinct and occupy the mid region of grey matter, Sp6-8 constitute the ventral horn, and Sp9 included almost exclusively MNs (Figure 2C). In thoracic section T4, we identified the visceral MN-containing lateral horn (LH). On the other hand, we were unable to define the dorsal nucleus of Clarke (expected in our T4 and L3 sections) based on enrichment of distinct Groups or consensus clusters. We expected that Sp4_mix4_Glut, which expresses genes like GDNF and POU4F1 (putative Clarke’s Column markers^25^), to define this nucleus; however, neither Sp4_mix4_Glut nor any other Group or consensus cluster was specific to this structure, suggesting that is defined by a combination of cell types or a subpopulation of one of the “mixed” Groups. Finally, we identified sparse (less than 10 cells per section) PITX2+ cholinergic interneurons in a column adjacent to the central canal (Figure S3) that were reported in mouse ^26^, we are not aware of prior work in primates that have described these cells.

We also noted that in the consensus UMAP projection (Figure 1C), Groups of the Glut-D Subclass fall into an apparent transcriptomic progression. To investigate whether this transcriptomic gradient corresponded to a spatial gradient, we used Partition-based graph abstraction (PAGA) to visualize whether the underlying cell type nearest neighbors graph mirrored laminar depth, as it does among cortical intratelencephalic-projecting types ^27^ (Figure 2D). Indeed, we found that 7 connected Groups followed a consistent relationship between transcriptome and depth (Figure 2D), and this relationship is more complex for mix1, TAC1, NMU TAC3, TAC3 NMU and SLIT2 dorsal glutamatergic Groups.

It has long been known that MNs differ in size across both regions of the spinal cord and across differently sized animals ^28,29^. To complement our spatial transcriptomic analyses, we performed immunofluorescence microscopy on adjacent macaque spinal cord sections to independently assess regional differences in neuronal density and soma size. The dorsal horn exhibited significantly higher cell density and smaller neuronal somata compared with the ventral horn (p < 0.05; Figures 2E and S4). We further quantified MN soma size using Feret diameter measurements from NeuN-stained cells, identifying the largest MNs in cervical and lumbar segments while thoracic and sacral had significantly smaller MNs (p < 0.05) (Figure 2F). Based on NeuN intensity and soma size, alpha and gamma MN subpopulations could be distinguished, consistent with prior reports ^30^. Notably, primate gamma MNs were significantly smaller than alpha MNs (p < 0.05; Figures S4 and S5), recapitulating size differences previously observed in rodent spinal cord ^31^. Comparison of primate MN sizes with reported measurements in human ventral horn lumbar cord revealed a consistent Feret diameter, on average, between species ^6,32^ (Figure 2F, Table S4).

Given the conservation of Group identities over 80 million years of evolution, yet striking differences in sensorimotor function and body size between mouse and primates, we looked for changes in Group abundances that may affect spinal cord circuits. We examined the relative composition of neuron types in segments in our droplet based sequencing and spatial data, and we observed an interesting Subclass-level allometric difference in the composition of neurons. We observed more cells from both GABA and glutamatergic dorsal and mid Groups relative to neurons from the ventral horn in a progression from mouse to macaque to human (Figure 2G).

### Comparative transcriptomics of consensus spinal cord types

We next looked at the conservation and divergence of the cell type expression in our consensus taxonomy. We find that many transcription factors (TFs) with Group-selective expression have well known roles in development and cell type identity across other regions of the central nervous system including LMX1B, MAF, SKOR2, LHX2, GBX1, BCL11A, PROX1, TFAP2A, and GATA3 (Figure 3A, Tables S5 and S6). We used the expressolog AUROC to measure expression pattern conservation of each gene across species, as well as the minimum Tau coefficient (see Methods) to measure cell type-specificity or ‘markerness’ of genes. We defined gene lists representing the intersection of markers and non-markers and conserved and non-conserved genes by dividing all 1:1 orthologs into quadrants defined by Tau=0.75 and AUROC=0.75. We found that conserved markers (high AUROC and high tau) had a significant enrichment in TFs (Fisher’s exact test p-value = 1.67e-11, Figure 3C). We found many well-known marker genes—like CHAT and TAC3—with high AUROC scores across species (Figure 3B).

**Figure 3.**
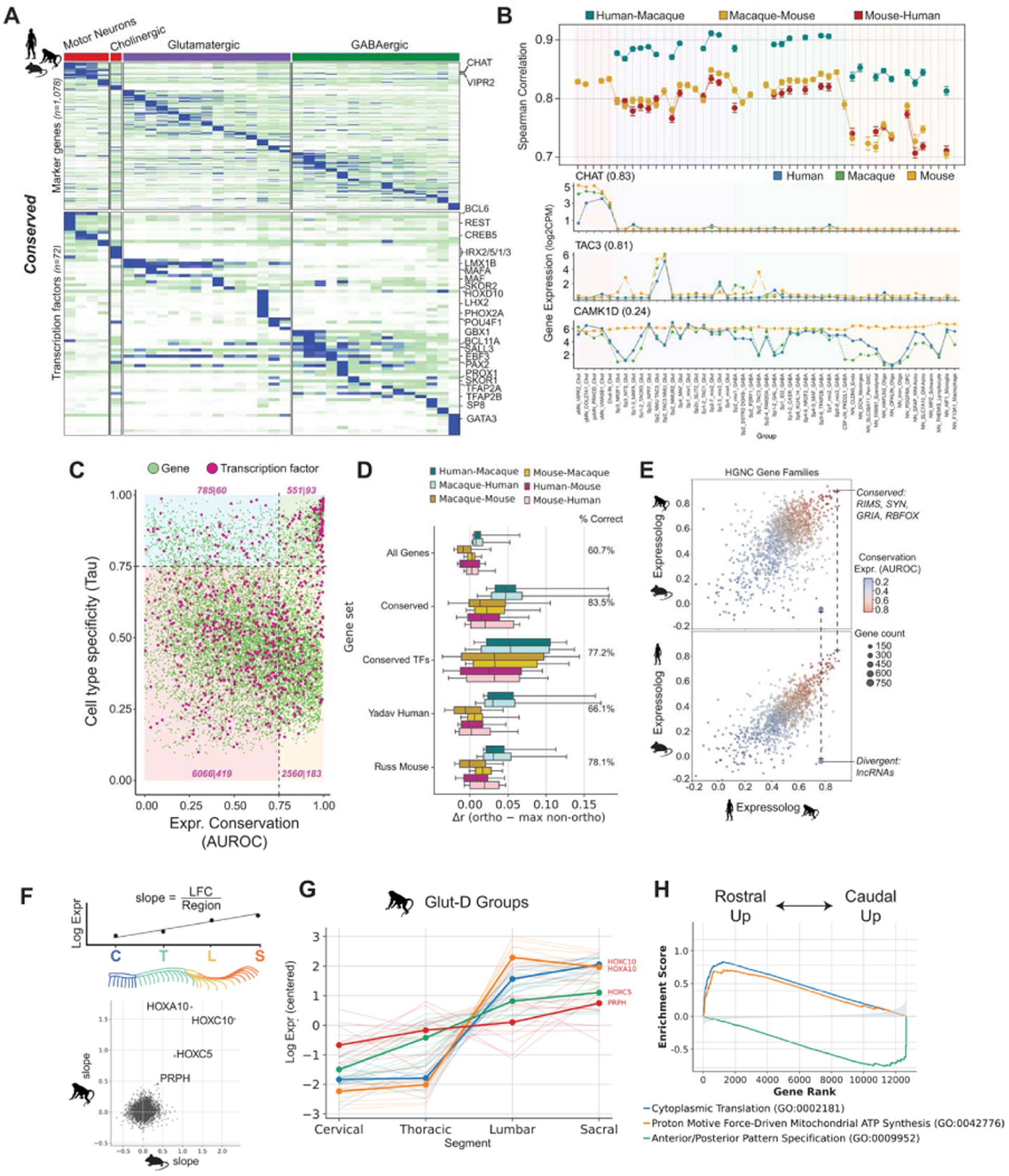
Comparative genomics of spinal cord cell types. **(A)** Heatmap of conserved marker genes and transcription factors across Groups in the consensus taxonomy (column-scaled mean expression across species). **(B)** Pairwise Spearman correlations of gene expression within well-supported Groups, error bars represent 95% bootstrap resampling confidence interval (500 resamplings of 100 cells, excluding species Groups with less than 100 cells). Below: Species-specific expression curves annotated by the mean expressolog correlation. **(C)** Ortholog conservation (x-axis) versus cell type specificity (y-axis). Quadrants show gene/TF counts: cons. markers (top-right), non-cons. markers (top-left), non-cons. non-markers (bottom-left), and cons. non-markers (bottom-right). **(D)** Self-projection accuracy of Groups and Subclasses using *MapMyCells* with variable gene sets. **(E)** Scatterplots of HGNC gene family expressolog scores between species pairs: Human vs. Macaque and Macaque vs. Mouse (top), Human vs. Macaque and Human vs. Mouse (bottom). Gene families of interest are labeled. Expressolog scores across hierarchical levels reveal persistent transcriptomic similarities. **(F)** Top: Schematic showing regression of regional Group mean to calculate slope in log fold change per region. Bottom: Scatterplot showing anterior-posterior gene expression mean slopes in the Glut-D Subclass, labeled genes have slopes with resampling p values less than 0.1 for more than 50% of Groups in the Glut-D Subclass. **(G)** Lineplot showing mean gene expression in each region of each Subclass for the significant genes from (F), with the highest rostral-caudal slope (light lines represent Groups within each Subclass). **(H)** Lineplot showing gseapy prerank directional enrichment of gene expression regression coefficients. Negative enrichment scores represent genes with increasing expression from rostral to caudal and vice versa.

We reasoned that with the high level of marker gene turnover seen across evolution, a marker set found in one species would be less effective in identifying that cell type in another species. We tested whether our conserved genes are features that provide improved separability in classifying cell type identity across species, compared to gene marker sets from any individual species (Table S6). We used different sets of genes to calculate correlations of each Group’s expression with its orthologous group in the other species, compared to the maximum correlation with non-orthologous Groups. We found that both conserved markers and conserved TFs yielded higher mapping correlations across species than gene sets derived from mouse or human taxonomies alone, and that 83.5% of Groups had the highest correlation with the correct orthologous Group using the conserved gene set, which outperformed gene sets found in human (66.1%) or mouse (78.1%; Figure 3D). Highly conserved gene families include RIMS, SYN, GRIA and RBFOX (Figure 3E), reinforcing deep conservation of presynaptic release, vesicle cycling, excitatory transmission and activity-dependent splicing programs. Primate-conserved and mouse-divergent genes (Figure S6) reflect diverse core cellular programs, including signaling (*CAMK1D* and *GRK4*) and extracellular-matrix biology (*LAMA1*), plus trafficking, structure, metabolism, and ion handling (Figures 3B and S3A). Divergent genes also include lncRNAs (Figure 3E) that may encode lineage-specific regulatory differences rather than protein-level changes, consistent with findings in neocortex^11^.

We summarized gene expression divergence from a cell type perspective using spearman correlation across all 1:1 orthologs (Figure 3B) or using DESeq2 differentially expressed genes (Figure S3). As expected, based on evolutionary distance, macaque and human cell type expression was far more similar to each other than either one was to mouse. In general, expression in neurons was more conserved than in non-neurons, with both Groups of astrocytes and oligodendrocytes appearing to be more diverged in primates compared to mouse by correlation, and with among the highest numbers of differentially expressed genes. We also saw that NN_PDGFRA_OPC was more conserved across species than the other glia, mirroring what is seen in the cerebral cortex ^33^. Interestingly, among neurons, we observed that dorsal Groups had lower correlations than ventral Groups, indicating a potential hotspot for evolutionary modulation (Figure 3B).

While consensus Groups included cells from across segments (Figure 1B), gene expression within Groups may still vary across the spinal cord. To characterize expression variation between segments, we regressed gene expression by the sampled region of the spinal cord and found that expression varied continuously along the rostral-caudal axis (Figure 3F). Looking at regression coefficients (calculated as the slope in log gene expression per region by assigning 0,1,2,3 to cervical, thoracic, lumbar and sacral, respectively), we found that across all developmentally endogenous spinal cord cell types (neurons, astrocytes, oligodendrocytes and ependymal cells), several caudal HOX genes like HOXA10 had the strongest directional expression change across segments, and genes with smaller regional expression differences included peripherin (PRPH) in dorsal Glut Groups and GABRQ in astrocytes (Figures 3G and S6). Across neurons, we tested the rankings of rostral-caudal slope magnitudes for gene ontology enrichments using gseapy^34^, identifying which types of genes tended to have more rostral-caudal or caudal-rostral expression. We found that metabolic genes related to protein translation and mitochondrial electron transport chain were more highly expressed in rostral cord, while HOX pattern specification-related gene sets were more highly expressed in caudal cord (Figure 3F-H).

### Comparative epigenomics of the mammalian spinal cord

To characterize gene regulatory evolution in the spinal cord, we generated cell type-resolved cis-regulatory element (cCRE) maps from single-nucleus ATAC-seq profiles in human, macaque, and mouse, identifying 0.75M, 1.25M, and 0.89M accessible cCREs, respectively. Sequence conservation was quantified using an updated 447-mammal Cactus multiple genome alignment from the Zoonomia Consortium ^35^ (see Methods). For comparative analyses, cCREs were classified by both sequence alignment and cross-species chromatin accessibility patterns (Figure 4A). Species-specific cCREs lacked ≥10% sequence alignment in any other profiled species, whereas sequence-conserved cCREs exhibited ≥50% alignment across all three species. Conservation of chromatin accessibility (epi-conservation) was quantified using an ‘accessolog’ metric, which measures similarity in Group-level accessibility patterns across sequence-conserved regions, analogous to expressologs ^36^.

**Figure 4.**
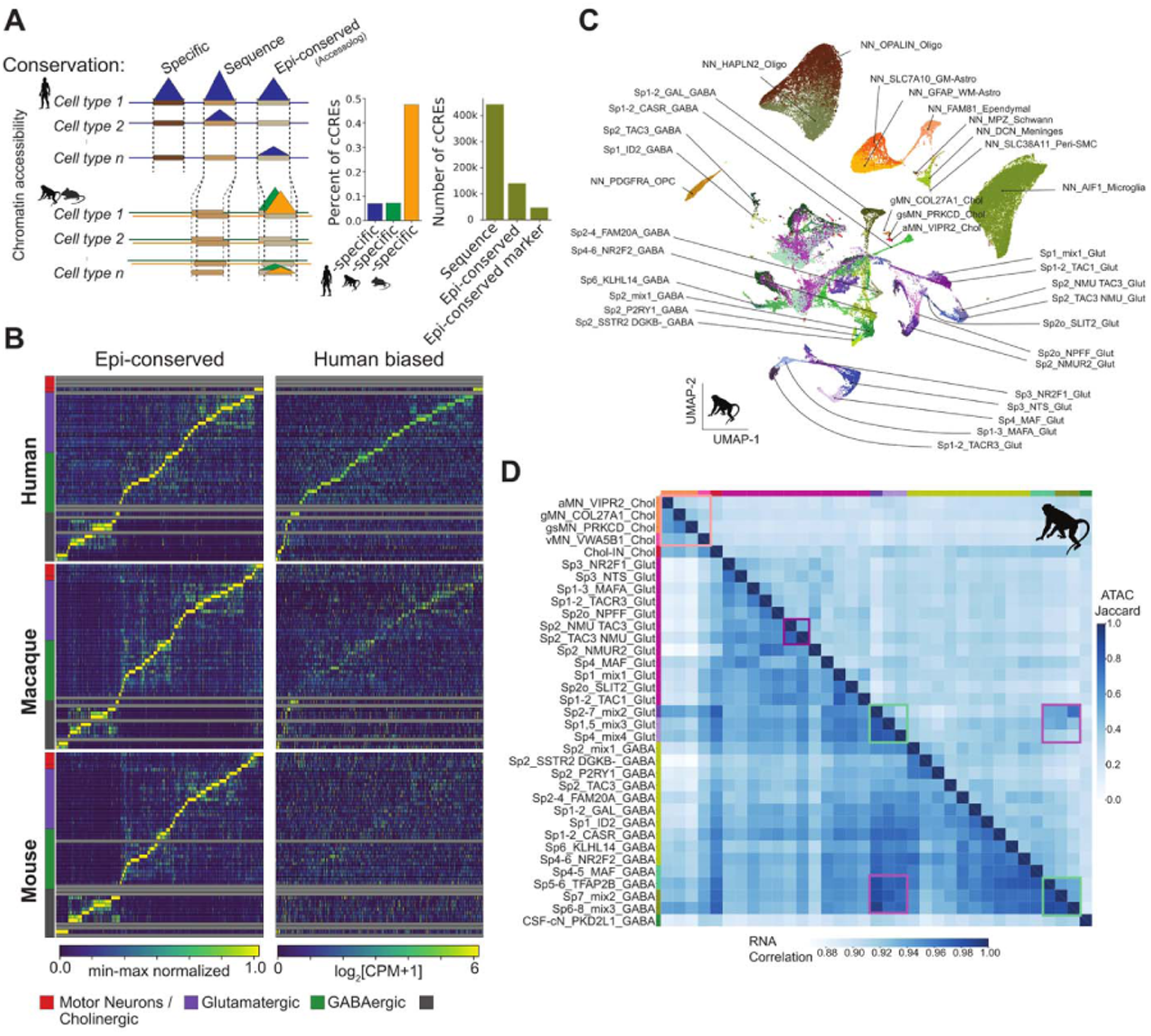
Comparative epigenomics identifies conserved and divergent Group cCREs. **(A)** Schematic of the levels of conservation for ATAC-seq peaks across species. Barplots show the number of peaks that fall into each category. **(B)** Heatmaps in which the highest specificity cCREs per Group are ordered accessibility for epi-conserved (top) and human-based cCREs (bottom). **(C)** Harmony batch-corrected UMAP projection of macaque single nucleus multiome ATAC profiles. **(D)** Heatmap showing the Pearson correlation (bottom left) and Jaccard index of binarized peak accessibility (top right) of macaque neuron Groups. Colored boxes highlight MNs (orange), similarity of ventral GABAergic and glutamatergic groups (purple) and ventral GABAergic types (green).

Using this framework, we observed substantial lineage-dependent differences in regulatory conservation. Approximately 7% of human and macaque cCREs were species-specific, whereas 43% of mouse cCREs lacked detectable conservation in primates, consistent with the greater evolutionary divergence between mouse and primates (Figure 4A). Across species, we identified 443,122 sequence-conserved cCREs spanning a broad range of cell-type specificities and enriched for promoter-proximal elements (Figure S4.1). Filtering sequence-conserved regions by high accessolog score further identified 140,120 epi-conserved cCREs with consistent accessibility across Groups, of which 47,609 were classified as epi-conserved marker cCREs based on high cell-type specificity and collectively spanning all Group terms (Figure 4A,B). Epi-conserved cCREs showed an enrichment of highly specific regions to neuronal Groups, including cholinergic as well as specialized GABAergic and glutamatergic neurons. In contrast, human-biased cCREs amounting to 0.3% of all cCREs were defined as sequence-conserved regions with elevated cell-type specificity uniquely in human and exhibited a stepwise reduction in specificity from human to macaque to mouse across all Group terms (Figure 4B). These human-biased elements were enriched in neuronal populations, indicating the emergence of lineage-specific regulatory programs layered upon a conserved chromatin accessibility architecture.

Highly variable chromatin accessibility profiles robustly separated nearly all Group terms defined in the RNA-based taxonomy, demonstrating that cCRE landscapes encode fine-grained spinal cord cell-type identity (Figure 4C). Dorsal GABAergic and glutamatergic populations, while transcriptionally closely related, segregated into distinct clusters within species-specific cCRE embeddings. Notably, TAC3-expressing glutamatergic neurons (Sp2_NMU_TAC3_Glut and Sp2_TAC3_NMU_Glut) were relatively similar in accessibility patterns across species compared to other types, consistent with shared regulatory programs (Figure 4C,D). MN Groups were well separated in cCRE space, with α MNs (aMN_VIPR2_Chol) forming a distinct cluster from the two γ MN populations (gMN_COL27A1_Chol and gsMN_PRKCD_Chol), demonstrating subtype-resolved regulatory organization within primate MNs (Figure 4C). In contrast, mid and ventral spinal cord GABAergic and glutamatergic populations exhibited shared co-accessibility patterns across Groups, which may suggest regulatory relationships not apparent in transcriptomic data, or be due to the heterogeneous nature of these groups (Figure 4C,D). These patterns suggest that chromatin accessibility captures aspects of regulatory organization that extend beyond steady-state gene expression, particularly in ventral spinal cord circuits.

### Sequence-based models reveal enhancer logic conservation across species

To investigate the sequence logic underlying Group-specific chromatin accessibility in the spinal cord, we trained three CREsted^37^ sequence models on the human, macaque, and mouse snATAC-seq datasets. Each model predicts chromatin accessibility directly from DNA sequence across all annotated spinal cord groups (Figure 5A). On held-out test sequences, the mouse, macaque, and human models achieved average Pearson correlations of 0.70, 0.73, and 0.51, respectively, between predicted and observed chromatin accessibility at peak regions on the genome (Figure S7), in line with significantly more cells profiled in macaque and mouse.

**Figure 5.**
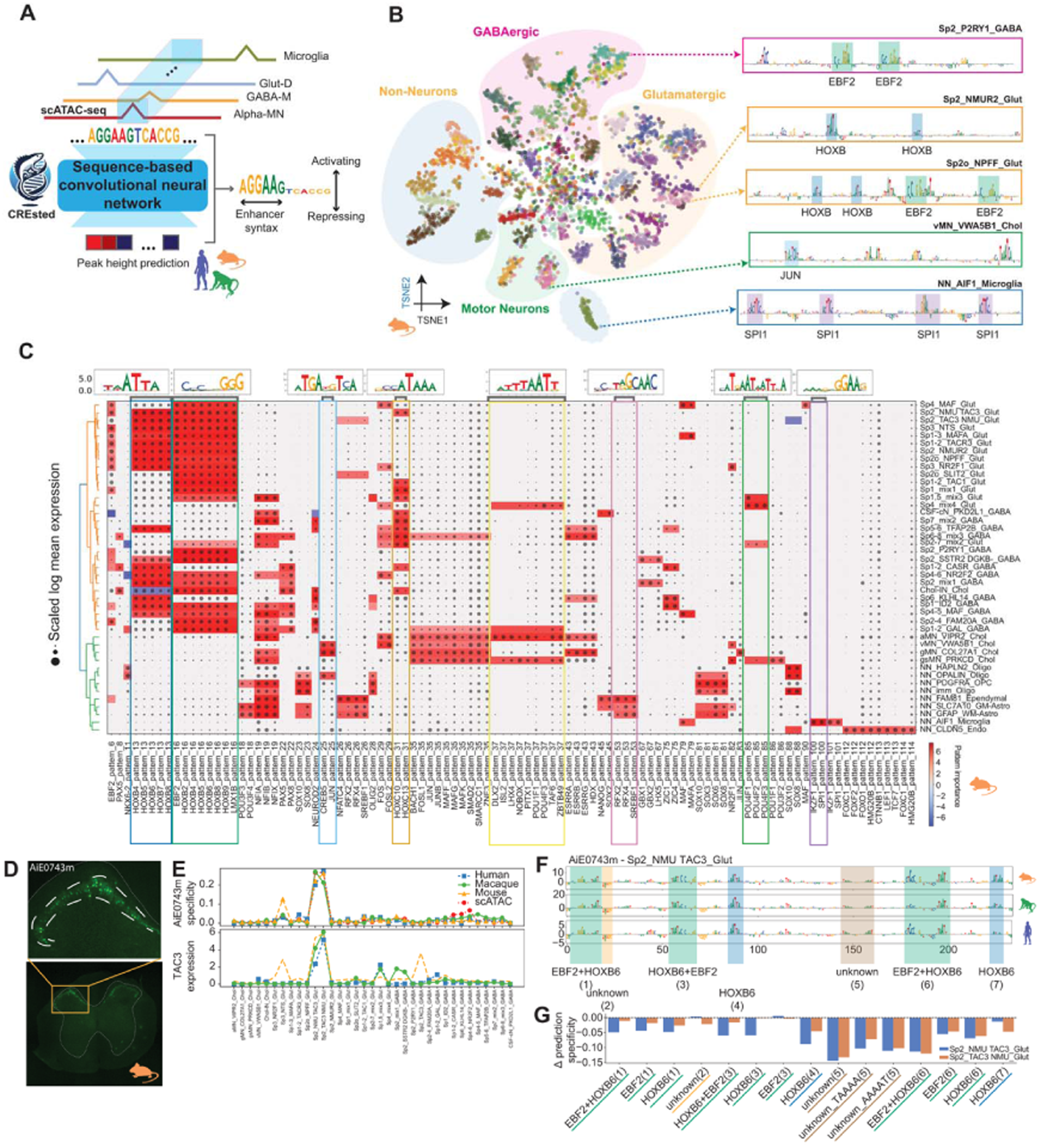
Sequence-based models reveal enhancer logic conservation across species. **(A)** Schematic of the training procedure for each species CRESted model and description of the per-base importance scores. **(B)** t-SNE of CREsted mouse model latent embeddings for the top 500 Group-specific cCREs colored by the Group to which each peak is specific. Contribution score plots of highly specific peaks for select cell types are displayed on the right with TF-motifs highlighted. **(C)** Cluster map of scaled mean log-normalized TF expression over Groups, indicated by dot size. Sequence patterns from CRESted matched to known TF motifs (columns) with mean importance per Group indicated by color. **(D)** Epifluorescence images of enhancer AiE0743m in mouse spinal cord sections. Fine dotted white line outlines grey matter, while heavier dashed white line outlines the approximate lamina 2. **(E)** Enhancer AiE0743m CRESted model predictions from each species and accessibility as measured from scATAC-seq for each Group. Expression pattern of TAC3 across groups and species. **(F)** Base-pair contribution scores for Enhancer AiE0743m for the species CRESted models with patterns from (C) highlighted in color. **(G)** Per-motif max contribution score values in dorsal glutamatergic Group terms relative to the NMU TAC3 and TAC3 NMU Groups, showing the change in overall peak specificity predicted associated with each motif.

To assess whether the CREsted models captured Group-specific regulatory logic, we extracted the internal embeddings of the mouse model for the top 500 most Group-specific peaks and visualized them after dimensionality reduction (Figures 5B and S7). These embeddings segregated sequences by broader neuronal Subclass and even by individual Group, indicating that the model encodes biologically meaningful enhancers. Applying the same analysis to the macaque and human datasets revealed consistent clustering patterns, and that ATAC prediction correlations between species were not as systematically different as transcriptome correlations (Figure S7), suggesting that enhancer logic is largely conserved across species (Figure S8). Scoring the top Group-specific regions from each species with all three models further supported this conservation with significant cross-species correlation in accessibility predictions (Figure S8)

Next, we examined the sequence features that drive these predictions. For each species, we identified motifs enriched in the 500 most specific regions per group and clustered them across all groups (Figures 5B and S8). We annotated motifs with candidate transcription factors supported by both snRNA-seq expression and motif-importance distributions (Figure 5C). Across species, EBF2 and HOXB motifs were broadly shared among neuronal classes, whereas DLX2/ISL2 motifs were most prominent in α-MNs, and POU4 motifs marked mid glutamatergic subtypes (Figure 5C).

Although the models displayed strong Group-specific predictive power, motif differences were more pronounced at the Class level than the Group level. This suggests that cell type specificity arises from combinatorial usage of a shared motif repertoire. To explore this further, we functionally tested several genomic elements targeting diverse cell types by AAV delivery in a mouse, examined their expression in spinal cord by reporter fluorescence and evaluated them using the CREsted models. Among these, we identified a mouse enhancer (AiE0743m was originally designed to target TACR3+ cells in telencephalon^38^) that drives reporter expression specifically in Sp2_TAC3 NMU/NMU TAC3_Glut cells within lamina 2 (Sp2) of the dorsal horn (Figure 5D). The enhancer showed Group-restricted accessibility in mouse snATAC-seq data and was predicted to be specific to the same types by all three CREsted models (Figure 5E). These Sp2_TAC3 NMU/NMU TAC3_Glut neurons across species express TAC3, a neuropeptide marker of neurons that have a critical role in mechanical itch processing^39^, linking enhancer activity to a conserved molecular identity.

To dissect the regulatory logic of enhancer AiE0743m, we computed nucleotide-level contribution scores across species (Figure 5F). All models converged on the same motif architecture, comprising multiple adjacent occurrences of EBF2-like and HOXB6-like sites, alongside two unannotated motifs. Systematic motif shuffling revealed that one of the unannotated motifs “unknown (5)” had the largest impact on predicted enhancer specificity (Figure 5G), suggesting that Group-specificity in glutamatergic neurons results from the combination of shared EBF2/HOXB6 elements and distinct, potentially novel, TF-binding sites with limited intersection with TF-motif databases. These analyses demonstrate that these models can accurately predict cell type-specific enhancer activity from sequence alone and uncover the combinatorial motif logic governing spinal cord regulatory specificity.

## DISCUSSION

In this study, we defined a cross-species, single-nucleus multiomic reference for the adult spinal cord that enables comparative and mechanistic analysis of spinal circuits across human, macaque, and mouse. By jointly constructing a transcriptomic consensus taxonomy, a cell type-resolved cis-regulatory landscape, and spatial maps, we define core organizational principles of spinal cord cell types and their association with anatomical architecture. We show that spinal cord cell types follow a deeply conserved molecular scaffold, with Classes, Subclasses, and Groups exhibiting clear conservation along the full length of the cord and across species. We show that this includes primate MNs, which have not been efficiently captured before, and that chromatin accessibility is distinct across Groups and reflects transcriptomic identity. Interestingly, we find that many of the conserved TFs marking cell types in spinal cord are well known as conserved core-identity TFs in other areas of the brain^40^. This finding reflects a strong developmental constraint, whereby the brain, as an ancient elaboration of the neural tube, reuses a shared and limited repertoire of ancestral transcriptional programs. Despite a conserved cell type architecture, we find large transcriptomic shifts across evolution, with thousands of differentially expressed genes between mouse and primate orthotypes. This turnover matches our findings that cross species profiling improves robustness of atlas mapping, as core identity genes are not apparent in one species alone. Together, this framework establishes a unified reference for mammalian spinal cord cell type organization and evolution.

Through our macaque spatial transcriptomic analyses, we showed not only that spinal cord cell types are approximately organized by traditional Rexed laminae, but also that the most fine-grained cell types (consensus clusters) had highly specific laminar distributions in cervical, thoracic and lumbar cord. Looking at the transcriptomic variation within Groups across the rostral-caudal extent of the cord, we expected to find many differentially expressed genes under HOX gene control. Surprisingly, we found that at the Subclass level, HOX genes were much more up-regulated in caudal cord than other genes, suggesting that this TF spatial gradient primarily controls transient developmental patterning rather than mature neuronal signatures, or that the effects vary by fine cell type. We also identified metabolism-associated genes that were up-regulated in the rostral cord, which is in agreement with studies measuring metabolic activity across the length of the human spinal cord in vivo ^41^. We also quantified the landscape of cell sizes and density from the cervical to sacral spinal cord, another cellular feature that will help bridge our results with other studies. Crucially, our consensus taxonomy now explicitly links known types with precise spatial distributions, making it possible to identify the expected spatial distribution of a cell type from its transcriptomic or epigenomic profile and facilitating integration of multimodal cellular properties across studies.

We observed a progressive enrichment of dorsal and intermediate neurons relative to ventral neurons in the profiled spinal cords from mouse to macaque to human. A prior MRI-based comparative study across mouse, rat, pig, monkey, and human did not report a strong corresponding shift in dorsal versus ventral horn area^2^. Together, these observations are more compatible with higher apparent dorsal neuronal packing in larger primates than with simple dorsal horn enlargement, but any evolutionary interpretation is tentative given only three profiled species and potential cross-species sampling effects. One plausible explanation is allometric scaling of sensory inflow with body surface area, whereas ventral motor circuit demands may scale more weakly. Alternatively, primate dexterity and locomotor differences could drive lineage-specific dorsal ventral allocation. Distinguishing these will require broader sampling across mammalian clades and body sizes.

Chromatin accessibility provided a complementary view of spinal cord cellular identity, revealing conserved fine-grained regulatory organization of cell Groups. By integrating sequence conservation with cell type-specific accessibility, we show that lineage-specific regulatory tuning is layered on a conserved cis-regulatory scaffold, particularly in neuronal populations. Mid and ventral interneurons exhibited shared co-accessibility patterns not evident from RNA alone, highlighting epigenomic state is a determinant of cellular identity. Sequence-based models trained independently in mouse, macaque, and human learned shared enhancer grammar that predicts cell type-specific accessibility across species, demonstrating that core regulatory logic is conserved. Importantly, enhancer specificity arises from combinatorial use of shared motifs rather than unique elements, providing a mechanistic basis for cross-species viral targeting and rational design of regulatory sequences that generalize from model organisms to human spinal cord cell types. Finally, from CREsted model analysis of the enhancer AiE0743m that is specific to the two Sp2_TAC3 NMU/NMU TAC3_Glut Groups, we found that the key motif driving specificity was an unknown sequence motif which was a TAAA…AAAT literal palindrome. This underscores the value of sequence models for prioritizing noncanonical regulatory elements for future experimental studies, including targeted mutagenesis and biochemical assays to identify the underlying DNA binding activity and mechanism.

Together, these results suggest that adult spinal cord cell types are built on a deeply conserved molecular and cis-regulatory program across species. Evolutionary differences include shifts in the relative abundances and gene regulatory programs of conserved cell populations, with more substantial remodeling of dorsal sensory circuits. By unifying transcriptomic, epigenomic, and spatial information across mouse, macaque, and human, this work provides a high-quality dataset and a consensus taxonomy for interpreting spinal cord cell types across studies, species, and modalities. More broadly, our analyses suggest a tractable path for bridging model organism experimentation to primate biology: conserved Group definitions enable direct alignment of circuit components, conserved cCRE grammar provides a mechanistic substrate for cross-species regulatory interpretation, and sequence models offer a scalable route to nominate and refine cell type-selective enhancers. As datasets expand in anatomical coverage, phylogenetic breadth, and functional annotation, this atlas should facilitate principled tests of how conserved spinal cord circuit motifs are modified to support species-specific behavior and how human-specific disease vulnerabilities may arise.

### Limitations of the study

Human neuronal sampling is limited, particularly for MNs, and integration with external human datasets can introduce batch structure. To mitigate this, we emphasize higher-confidence, cross-species Group-level correspondences supported by within-study replication and concordant transcriptomic and epigenomic evidence. Spatial data was limited to macaque, and laminar boundaries were not consistently recovered by automated approaches due to curvature and overlap of Rexed laminae. We therefore defined laminar domains manually from MERFISH cell-type distributions and cross-checked these assignments with adjacent-section histology. Finally, a subset of ventral populations remains aggregated as “mix” Groups in the consensus taxonomy. We retain these as explicit placeholders with marker and anatomical context and refer readers to the companion work (Cano Gomez et al. 2026) for higher-resolution subdivision.

## RESOURCE AVAILABILITY

### Lead Contact

Further information and requests for resources and reagents should be directed to and will be fulfilled by the lead contact, Trygve E. Bakken.

### Materials Availability

This study did not generate new unique reagents.

### Data and Code Availability

- Sequencing data and enhancer validation data used in this project are all from previous studies and are publicly available. Accession numbers and links to the datasets are listed in the key resources table.
- All original code has been deposited at GitHub and is publicly available as of the date of publication. GitHub URLs and DOIs are listed in the key resources table.
- Any additional information required to reanalyze the data reported in this paper is available from the lead contact upon request.

## ACKNOWLEDGMENTS

This publication was supported by and coordinated through the BRAIN Initiative Cell Atlas Network (BICAN) (https://braininitiative.nih.gov/research/tools-and-technologies-brain-cells-and-circuits/brain-initiative-cell-atlas-network). This work was funded by the Allen Institute for Brain Science and by the National Institutes of Health U24MH130918 (PLR), U24MH130919 (PLR), and UM1MH130981 (AA, AB, ABC, AO, AR, AT, BT, CH, CR, CS, CTJvV, DH, DM, DN, EK, EL, ESL, HZ, IK, JA, JG, JN, JTT, JW, KAS, KF, LN, LS, MH, MR, MTS, MW, ND, NJJ, NP, NVC, NVS, NXM, PO, RDH, RF, SB, SCS, SD, SDH, SRK, SY, TC, TEB, TLD, WH, YF, YG, ZCJ, ZRL, ZY). Research Foundation Flanders (FWO) PhD fellowship 1SH6J24N & V428025N (NK). The authors thank the founder of the Allen Institute, Paul G. Allen, for his vision, encouragement and support.

## AUTHOR CONTRIBUTIONS

Tissue acquisition: CH, EL, LS, MR, MTS, RDH, TLD

Sample preparation and multiomic data generation: AT, CR, CS, DH, HP, KAS, LC, NP, NVS, RDH, RF, TC, WH, KF, ET

Spatial transcriptomic data generation: AA, AO, AR, DM, JA, JN, JW, MH, NVC, NXM, PO, SB, SCS, SDH, SRK, ZCJ

Viral tool testing: BT, DN, EK, ND, SY, TLD

Data archive / Infrastructure: AB, ABC, JG, KAS, LN, PLR

Data analysis: BT, CTJvV, IK, JG, KAS, MH, MTS, NJJ, NK, SCS, SD, TEB, TLD, YF, YG, ZY

Data interpretation: A-HP, ADG, AP, BT, CTJvV, ESL, HL, IK, JAB, JTT, KF, MH, MJL, MN, MTS, MW, NJJ, NK, OG, RDH, RS, SCS, SD, TEB, TLD, YF, YG, ZRL, ZY

Writing manuscript: ESL, IK, MTS, NJJ, NK, TEB Funding acquisition: ESL, HZ, RDH, TEB, ZY

## DECLARATION OF GENERATIVE AI AND AI-ASSISTED TECHNOLOGIES

During the preparation of this work, the authors used OpenAI ChatGPT-5 to refine the main text by improving conciseness and readability, and to assist in identifying potentially relevant citations. After using this tool, the authors reviewed and edited the content as needed and take full responsibility for the final content of the publication.

## STAR METHODS

### Macaque Tissue Specimens

Macaque spinal cord tissue samples were obtained from the University of Washington National Primate Resource Center tissue distribution program under a protocol approved by the University of Washington Institutional Animal Care and Use Committee. Immediately after euthanasia, macaque spinal cords were removed and transported to the Allen Institute in artificial cerebral spinal fluid equilibrated with 95% O2 and 5% CO2. Upon arrival at the Allen Institute, spinal cord samples were subdivided, marked with black ink to maintain anatomical orientation, and immediately flash frozen in a bath of dry ice and isopentane. Frozen spinal cords were transferred to vacuum sealed bags and maintained at −80°C until the time of tissue dissection. Dissection was carried out on a custom cold table maintained at −20°C. Spinal cord samples were removed from the −80°C freezer and briefly warmed to −20°C, after which they were manually subdivided into ∼2mm thick slabs using a standard razor blade. For all samples, excess white matter was removed and to aid in enrichment of neuronal populations. For a subset of samples, the ventral horn of the spinal cord was dissected using a dry ice cooled fine scalpel and processed for nuclei isolation separately.

### Human Tissue Specimens

Tissue collection was performed in accordance with the provisions of the United States Uniform Anatomical Gift Act of 2006 described in the California Health and Safety Code section 7150 (effective 1 January 2008) and other applicable state and federal laws and regulations. The Western Institutional Review Board reviewed the use of these tissues and determined that they did not constitute human subject research requiring institutional review board (IRB) review. Male and female individuals 18 to 68 years of age were considered for inclusion in the study. Donor screening involved a review of basic clinical metadata and serological testing for infectious diseases. Standard tissue quality metrics (RNA integrity evaluation [RIN], pH, tissue slab quality) were assessed for all donors and a RIN value of ≥7.0 was required for inclusion in the study. Donor tissues for the current study were obtained from four donors (3 male, 1 female, ages 27-42 years) via the University of Washington BioRepository and Integrated Neuropathology (BRaIN) laboratory. A rapid autopsy was performed on each donor where the brain was removed, bisected along the midline, and each hemisphere was embedded in alginate. Cerebral hemisphere and cerebellum tissue slabs were generated separately from the brainstem and spinal cord. The brainstem was slabbed by hand using an autopsy knife to grossly separate the midbrain, pons, and medulla including the most anterior part of the cervical spinal cord, each of which were frozen as separate samples in a bath of dry ice and isopentane. In one donor, the full spinal cord was removed at the time of autopsy and subdivided into ∼5 cm long segments prior to freezing in dry ice and isopentane. All tissue slabs were transferred to vacuum sealed bags and stored at −80°C. Tissues were transferred to the Allen Institute on dry ice and were stored at −80°C until the time of dissection. Tissue dissections were conducted on the custom cold table described above at a constant temperature of −20°C. Subdivision of spinal cord blocks and dissection of ROIs was done as described for macaque.

### NIMP Integration

In collaboration with the BICAN consortium, fresh tissue images captured for all spinal cord specimens were uploaded to the Neuroanatomy-anchored Information Management Platform (NIMP) (Tao et al., co-submitted)^42^. Slab photos were digitally cropped and then assigned to dissection-specific requests in NIMP, which allowed dissectors to draw dissection plans and assign anatomical pins linking drawn regions of interest (ROIs) with neuroanatomical structures defined in our HOMBA ontology (https://alleninstitute.github.io/CCF-MAP/docs/HOMBA_ontology_v1.html) (Ding et al., co-submitted)^43^.

### Sample Processing for 10x Genomics Multiome

Single-nucleus suspensions were generated using a previously described standard procedure (https://dx.doi.org/10.17504/protocols.io.5qpvok1exl4o/v2). Briefly, after tissue homogenization, isolated nuclei were stained with primary antibodies against NeuN (FCMAB317PE, 1:100 dilution, Sigma Aldrich) to label neuronal nuclei and OLIG2 (Abcam ab225099, 1:400 dilution) to distinguish non-neuronal populations. Nucleus samples were analyzed using a BD FACSAria II or FACSAria Fusion flow cytometer (software BD Diva v.8.0, BD Biosciences); nuclei were sorted using a standard gating strategy to exclude multiplets. A defined mixture of neuronal (70% from the NeuN+ gate) and non-neuronal (20% from the OLIG2− and 10% from the OLIG2+ gate) nuclei was sorted for each sample. After sorting, nuclei from all sorted populations were combined and concentrated by centrifugation. Chip loading and postprocessing to generate sequencing libraries were done with the Chromium Next GEM Single Cell Multiome Gene Expression kit according to the manufacturer’s guidelines. Nuclei concentration was calculated either manually using a disposable hemocytometer (DHC-NO1, INCYTO) or using the NC3000 NucleoCounter. All 10X libraries were sequenced according to the manufacturer’s specifications on a NovaSeq 6000 using either a NovaSeq X or S4 flow cell. Reads were demultiplexed to FASTQ files using BCL Convert (v.4.2.7) for libraries run on NovaSeq X flow cells and bcl2fastq (v.2-20-0) for libraries run on S4 flow cells. Reads from snRNA-seq libraries were mapped to the 10X Genomics official human reference (GRCh38-2020-A); UMIs per gene were counted using the Cell Ranger (v.6.1.1) pipeline with the --include--introns parameter included. Reads from the snATAC–seq and snMultiome libraries were mapped to the same reference using Cell Ranger ATAC (v.2.0.0) and Cell Ranger Arc (v.2.0.0) pipelines, respectively, with default parameters.

### Quality Control of snRNA-seq data

We performed multi-stage quality control (QC) across species when constructing the consensus taxonomy. For each species, we applied basic cell-level QC to remove low quality profiles. Cells with fewer than 1,000 detected genes or 2,000 UMIs were excluded, and cells with abnormally high gene or UMI counts were removed as well. After an initial round of clustering and label-transfer annotation, we performed a label-aware QC pass at the cluster-level. Clusters with low median gene or UMI counts, elevated doublet scores, or highly heterogeneous broad annotation profiles were removed. In addition, once major neuronal and non-neuronal identities were established, we applied a stricter neuronal filter by excluding neurons with <2,000 detected genes. Minor variations in thresholds were applied across species to account for differences in data characteristics.

Next, high-quality cells from all species were aligned in a shared embedding space. Clusters showing poor cross-species alignment were manually reviewed based on multiple criteria, including within-species embedding structure, species-mixing patterns in the aligned space, and expression patterns of marker genes. Clusters that failed alignment criteria were excluded. Finally, evidence from spatial transcriptomics and additional multiomic assays was used to further corroborate cluster identity, assess anatomical consistency, and refine cross-species of QC decisions.

### Clustering of snRNA-seq data

Clustering was performed using a Python implementation of the scrattch.hicat iterative clustering package (https://alleninstitute.github.io/scrattch.hicat/). To minimize donor and species-specific variation prior to clustering, we generated a single donor and species-corrected latent representation (via scVI) and used this corrected embedding throughout the entire iterative procedure. No additional donor-level adjustment was performed during downstream rounds of subdivision.

We applied the iter_clust automatic top-down clustering function which recursively partitions cells into increasingly fine transcriptomic groups. This algorithm identifies new clusters only when all resulting subclusters are mutually separable by stringent differential gene expression (DGE) criteria. Following Yao et al. (2023)^8^, we used parameters optimized for detecting small but transcriptionally distinct populations:

- q1.th = 0.5 (minimum fraction of expressing cells in the candidate foreground cluster),
- q.diff.th = 0.7 (minimum normalized expression-fraction difference distinguishing foreground and background),
- de.score.th = 150 (minimum cumulative differential-expression score), and
- min.cells = 4

These criteria ensure that any pair of final clusters is separated by at least 8 binary DEGs. Each DEG’s contribution to de.score is capped at 20, so a minimum of eight genes is required to exceed the threshold of 150. Binary DEGs were defined as genes expressed in ≥40% of cells in the foreground cluster, with |log2 fold-change| > 1, adjusted P < 0.01, and a foreground-normalized expression-fraction difference > 0.7.

### Annotation transfer with MapMyCells

Transcriptomic annotations were transferred onto the HMBA Spinal Cord dataset using MapMyCells (RRID:SCR_024672) implemented in the Python package cell_type_mapper (*AllenInstitute/cell_type_mapper*). Briefly, a curated reference atlas with taxonomic labels was used as the training reference, and HMBA spinal cord cells were treated as the query. MapMyCells then computed gene-level similarity scores and performed both correlation and hierarchical label transfer, assigning each HMBA cell the most likely reference label at multiple taxonomic levels (e.g., Class, Subclass, Group). The method returns both discrete labels and confidence scores, which we used to filter low-confidence mappings and to refine downstream consensus annotations.

### Replicability of Group-terms across species with MetaNeighbor

Group-level replicability across species was evaluated using MetaNeighbor^44^, which quantifies how well cell type labels can be recovered using cross-dataset classification. Expression matrices were restricted to the conserved marker genes, and transcriptomic Groups were used as labels. For one-vs-best replicability, MetaNeighbor identifies, for each target and reference species pair, the two closest matching transcriptomic Groups and computes an AUROC based on the voting results, comparing cells from the closest match to cells from the second closest match. The resulting AUROC values reflect how distinctly each Group can be identified across datasets in a local context. High one-vs-best AUROC scores (>0.7) indicate strong Group replicability, whereas lower values highlight Groups with weaker cross-dataset correspondence.

### MERSCOPE data generation and processing

Post-processing of the macaque MERSCOPE data used custom tools in a pipeline consistent with Hewitt et al. 2025^45^. Because the MERSCOPE platform cell segmentation tools failed to recognize MNs, we chose to re-segment with a custom segmentation pipeline that showed improved data quality on the basal ganglia^45^. The software is available on Github (https://github.com/AllenInstitute/spots-in-space/tree/main/sis). In brief, re-segmentation of the macaque MERSCOPE data was performed using a custom 3D Cellpose2 model trained on our MERSCOPE data using a human-in-the-loop approach, ensuring that both MNs and other cells were segmented. Two channels were used: the DAPI image acquired during MERSCOPE acquisition, and an image created from the total mRNA signal of the MERSCOPE data to serve as a cytosol stain. However, we note that even with this custom approach, whole MNs are not segmented as single cells but instead are often into multiple segmented cells.

### MERSCOPE data taxonomy mapping

Cells were mapped to our consensus spinal cord snRNA-seq reference dataset using MapMyCells’ (RRID:SCR_024672) hierarchical mapping algorithm with bootstrapping (100 iterations, bootstrap factor 0.5). We used all taxonomy levels, Class, Subclass, Group and consensus_cluster. We filtered cells with less or equal to 10 gene counts, or cells with an average bootstrap Group correlation < 0.1. We also used cellxgene to manually segment spinal cord grey matter, and approximate lamination using Group mappings.

### Quantification of neuron sizes and densities

Motor neurons were identified and quantified using NeuN image volumes in Neuroglancer (v2.41.2), a web-based visualization tool for large-scale volumetric data. We used a Neuroglancer build that includes the polyline annotation type introduced in the merged pull request #769, enabling manual tracing of motor neuron outlines across multi-scale, high-resolution views. Motor neurons were manually segmented based on their large soma size and characteristic morphology in the ventral horn, in contrast to the smaller neuronal somata observed in the dorsal horn. γ-motor neurons (γ-MNs) were further distinguished by their absence of NeuN labeling or markedly reduced NeuN signal intensity relative to α-motor neurons.

Annotated polylines were exported as JSON files and analyzed in Python 3.13 using Shapely to construct closed polygons and compute morphometric features. Motor neuron soma size was quantified using the Feret diameter, defined as the maximum pairwise Euclidean distance between vertices of the polygon’s convex hull. Measurements were aggregated across α- and γ-motor neurons, merged across donors, and compared across spinal segments using Welch’s two-sample t-tests.

Whole section neuronal soma size and local spatial density were quantified on NeuN stains from high-resolution segmentation masks generated using a Cellpose (v 4.0.6) model trained via reinforcement learning and subsequently refined by manual curation for each section. All measurements were converted from pixel units to micrometers using section-specific pixel-to-micron scaling factors.Size was quantified using the Feret diameter, or caliper diameter, defined as the maximum pairwise Euclidean distance between all pixels belonging to a neuron. To characterize dorsoventral trends in soma size, neurons were ordered along the dorsoventral axis and smoothed using a centered rolling median window.

Neuron spatial density was quantified using an outline-based neighborhood analysis. Neuron boundaries were extracted from segmentation masks, converted to micrometer coordinates, and indexed using a KD-tree. For each neuron, neighboring neurons were identified as those with boundary pixels within a fixed radius (100 µm), and the number of unique neighboring neuron identities was recorded as the local neighborhood density. Neighborhood density values were analyzed as a function of dorsoventral position using the same spatial normalization and rolling median smoothing as for soma size.

### ATAC-seq processing, quality control, and annotation

Single-nucleus ATAC-seq data from 10x Genomics multiome libraries were processed with snapatac2 ^46^ (version 2.8). For each sample, we imported fragment files into an on-disk AnnData object using *snap.pp.import_fragments* with appropriate reference (fasta, GTF, and chromosome sizes). We restricted analysis to nuclei that were also present in the matched RNA-seq dataset, enabling direct transfer of transcriptomic labels (Class, Subclass, Group, and cluster ID) from the RNA to the ATAC modality on a per-nucleus basis via shared barcodes.

Quality control was performed in snapatac2 using TSS enrichment, nucleosome signal, and fragment counts as key metrics. TSS enrichment was computed with *snap.metrics.tsse*, and nuclei were retained if they had TSS enrichment ≥ 5 and ≥ 1,000 fragments (*snap.pp.filter_cells(min_tsse=5.0, min_counts=1000*).

To generate bigWigs, we concatenated all high-quality nuclei into a single snapatac2 dataset and computed coverage grouped by transcriptomic Group. TSS regions were defined using GENCODE annotations: all “transcript” entries in the GTF were converted into ±100 bp windows around the transcription start site, and overlapping windows were collapsed to a non-redundant set of TSS intervals. These TSS intervals were supplied to *snap.ex.export_coverage* and coverage was exported in 10 bp bins using insertion counts and CPM scaled by TSS reads, such that each bin value equaled *(reads in bin / total reads in TSS intervals) × 10*_. This procedure yields cross-sample and cross-Group bigWigs normalized to TSS-proximal signal.

Peak calling was performed in snapatac2 via MACS3 using default parameters, applied per Group (*snap.tl.macs3*). Group-specific peak sets were then merged across Groups and chromosomes using *snap.tl.merge_peaks* to generate a unified, non-redundant peak catalog. A peak-by-cell count matrix was built from this merged peak set with *snap.pp.make_peak_matrix*, and the top 250,000 most variable peaks were selected as features (*snap.pp.select_features*).

For dimensionality reduction and donor correction, we computed a spectral embedding (*snap.tl.spectral*) and UMAP visualization (*snap.tl.umap*) on the peak matrix. Donor identities were mapped to each nucleus by parsing sample IDs from cell barcodes and joining to external metadata. To control for donor-specific effects, we applied Harmony batch correction on the spectral embedding (*snap.pp.harmony(batch=“donor”)*) and computed UMAP using the Harmony-corrected representation. This donor-corrected latent space was used for all downstream ATAC-based visualizations.

### Expressologs

We quantified cross-species conservation of gene expression patterns, expressologs ^36^, using pseudobulk mean expression matrices for human, macaque, and mouse for each annotation level (Class, Subclass, Group) and shared set of 1:1 orthologous genes. Cell type labels were harmonized by taking the union of all observed cell types; for any missing cell type in a given species, a zero-filled row was added to ensure that all matrices were aligned and identically ordered. For each pairwise comparison, we designated one species as the reference and the other as the query. Then for every gene in the shared ortholog set, we computed the Pearson correlation between its pseudobulk expression profile across matched cell types in the reference and query species. We also recorded the standard deviation of each gene’s expression profile in each species as a measure of variability. This process produced a symmetric set of gene-wise expression correlations for all primate species pairs (reference→query and query→reference).

To derive expressolog scores for 1:1 orthologous gene pairs, we standardized these pairwise correlations using a rank-based AUROC procedure.

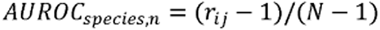

where N is the total number of genes, and r_ij_ is the rank of the Pearson correlation of gene i with its 1:1 ortholog gene j relative to all other (N□−□1) genes in the second species.

For each gene, we computed the median cross-species expressolog score across ortholog comparisons.

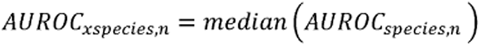

The expressolog scores were also calculated across taxonomy levels using the same formula and captures the extent to which gene expression variation is shared across species.

### Cell type specificity metric

We quantified gene-level cell type specificity across species using a Tau-based meta-analysis of snRNA-seq expression profiles^47^. For each species (human, macaque, mouse), we used AnnData objects containing donor-resolved cell-by-gene count matrices and consensus annotations (“Class”, “Subclass”, “Group”). To obtain robust cell type-level estimates while accounting for donor heterogeneity, we performed repeated donor subsampling: for each species, in each iteration we randomly selected a subset of donors (number of donors = max[2, ceil(total_donors/3)]) and restricted the analysis to cells from those donors. Within each subsample, we aggregated expression per annotation level using scanpy utilities to compute mean expression for each gene in each cluster. Genes with mean expression > 1 in at least one cell type were retained, and the resulting gene × cell type matrix was used to compute Tau scores.

Tau was defined for each gene as:

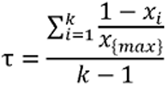

where (x_i) is the mean expression in cluster (i), (x_{\max}) is the maximum mean expression across the (k) clusters, and genes with zero maximal expression were assigned Tau = 0. This procedure was repeated for 10 random donor subsamples, yielding a matrix of Tau values (one column per subsample) for every gene. We then summarized the mean Tau across subsamples (“mean_tau”) and used this value for downstream analysis.

### Cactus sequence alignment

To prepare the 447-way Cactus alignment for use with the specific genome assemblies used with our genomics data, the replace utility of the Cactus-update-prepare was used for the macaque and marmoset genomes. Cross-species orthologs of ATAC-seq peaks were identified using the Zoonomia 447-mammal Cactus HAL alignment ^48^. For each species, merged peak BED files were projected across the Cactus alignment using HALPER^49^ and halLiftover (*pfenninglab/halLiftover-postprocessing*), which identify orthologous intervals in other genomes while enforcing alignment-quality constraints.

We used the following HALPER parameters to ensure high-confidence mappings:

- MIN_LEN = 50: minimum aligned length required for an orthologous interval
- PROTECT_DIST = 10: distance flanking the query interval preserved during alignment to avoid edge truncation
- MAX_FRAC = 2: maximum allowed fractional expansion of the orthologous region relative to the input peak (i.e., up to 2× peak length)

Peak BED files were mapped across all species in the 447-way HAL, and HALPER postprocessing steps were applied to merge, filter, and classify orthologous intervals. The resulting tables provided, for each input peak, the set of confidently aligned orthologs across species and ancestral nodes, enabling downstream evolutionary and conservation analyses.

### Definitions of ortholog and species-specific alignments

Using the output of HALPER on the 447-way Cactus alignment, we defined orthologous cCREs among the three profiled primate species (human, macaque, mouse) in a human-anchored framework. We started from merged human ATAC cCREs and HALPER narrowPeak files that align macaque and mouse peaks into human (hg38) coordinates. Human peaks were given unique genomic IDs (chr:start–end) and converted to PyRanges; for each non-human species, we filtered HALPER alignments to retain only intervals with ≥250 bp aligned to the human peak (∼50% of a 500-bp cCRE), corrected HALPER-set peak IDs. We then intersected species-specific HALPER tracks with the human cCRE set to assign, for each human region, the corresponding macaque and mouse peak IDs and aligned human coordinates. This produced a human-anchored liftover table from which we classified species-specific cCREs as those lacking matches in both other species, and orthologous cCREs as those with aligned peaks in all three species.

### cCRE annotation

We annotated ATAC cCRE for overlap with transposable elements (TEs) using species-specific RepeatMasker tracks. For each species, merged peak BED files were converted to PyRanges objects and given a unique peak ID (chr:start-end). Species-appropriate RepeatMasker annotations were loaded in UCSC rmsk schema (hg38, rheMac10, mm10). Repeat records were converted to PyRanges and intersected with peaks using a genomic join. For each peak-TE overlap, we retained the repeat name (repName), class (repClass), and family (repFamily), and required a minimum overlap of 250 bp (∼50% of a 500-bp peak); overlaps below this threshold were treated as non-TE. cCREs lacking qualifying overlaps were labeled with False and −1 for TE and repeat fields. The resulting TE-annotated peak tables were saved per species for downstream analyses.

We annotated cCREs for promoter overlap using species-specific GENCODE gene models. For each species, merged cCRE coordinates were converted to PyRanges objects with unique region identifiers. Promoter intervals were derived from GTF annotations by extracting protein-coding gene entries, computing strand-aware transcription start sites (TSS), and expanding these to asymmetric promoter windows (–1 kb / +500 bp on the positive strand; –500 bp / +1 kb on the negative strand). Species-specific chromosome aliases were applied for macaque and mouse. cCREs were intersected with promoter intervals using genomic joins, and each region was assigned a binary promoter label based on whether it overlapped any promoter interval. For each species, promoter-annotated cCRE tables were exported for downstream analyses.

We quantified evolutionary conservation for human and cross-species orthologous cCREs using phyloP 100-way vertebrate scores. Human cCREs along with macaque and mouse orthologous cCREs in hg38 coordinates were combined into a unified set of species-cCRE tables. For each cCRE, we retrieved base-resolution phyloP scores from the hg38 phyloP100way bigWig using pyBigWig. We queried each cCREs chromosome, start, and end coordinates; extracted the corresponding phyloP vector; removed missing values arising from unaligned or low-coverage regions; and computed the mean phyloP score cCREs with no valid phyloP positions were assigned NaN.

### Identification of cell type-specific cCREs

The Gini coefficient provides a single-value summary of how unevenly a cCRE’s accessibility is distributed across cell types. For each cCRE, we treat its pseudobulk accessibility across k cell types as a vector *x* = (*x*_1_, …, *x_k_*) *x* = (*x*_1_, …, *x_k_*).The Gini coefficient measures inequality in this vector, taking a value of 0 when accessibility is evenly distributed across all cell types and increasing toward 1 as accessibility becomes concentrated in a single cell type. Formally, the Gini coefficient is computed by sorting accessibility values in ascending order, then evaluating for each cCRE:

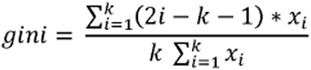

Thus, cCREs with broad accessibility have low Gini scores, whereas cell type-restricted or highly specific cCREs yield high Gini values. In our pipeline, we report for each cCRE the maximum Gini value across cell types (assigned to the cell type with the highest accessibility), which serves as the cCRE’s overall specificity score.

### Conservation of chromatin accessibility

To integrate sequence conservation with chromatin-state conservation, we defined **accessologs**, the chromatin accessibility analogue of expressologs. Accessologs quantify whether a sequence-conserved cCRE retains a similar pattern of accessibility across homologous cell types in another species. Using ortholog regions derived from the Cactus multi-species alignment, we computed pseudobulk accessibility for each cCRE across the consensus spinal cord cell types. For every pair of species, we calculated the Pearson correlation between the accessibility profile across matched cell types in the reference and query species. We then rank-standardized the similarity of the true orthologous region relative to all possible region pairs, yielding an **AUROC-based accessolog score**. High accessolog scores indicate that a conserved sequence element also preserves coordinated accessibility patterns across species, whereas low scores reflect divergence in regulatory activity despite sequence alignment.

### DNA sequence model training

We trained three CREsted ^37^ models that predict cell type-specific accessibility from sequence using the mouse, macaque, and human snATAC-seq data. The aggregated peak signal was additionally scaled over groups using *crested.pp.normalize_peaks()* on the top 3% of peaks per group. Training, validation, and test sets were split by chromosome: chr10 for validation set, chr18 for test set, the remaining for training. For all models, we used the default peak regression training strategy and configuration. We pretrained our models on all consensus peaks, and then fine-tuned to variably accessible regions (defined by *crested.pp.filter_regions_on_specificity()*). Fine-tuning was done with a lower learning rate (1e-4) than was used during the pre-training.

### Sequence embeddings

We computed a two-dimensional t-SNE embedding of regions based on sequence representations extracted from the mouse CREsted model. For each species, we took the top 100 most specific regions that were selected using *crested.pp.sort_and_filter_regions_on_specificity()* with method=”proportion”. We then calculated embeddings from the *global_average_pooling_1d* layer of the mouse CREsted model via *crested.tl.extract_layer_embeddings*(), and the resulting matrices were concatenated across species. The combined embedding matrix was reduced to two dimensions using t-SNE (sklearn.manifold.TSNE, n_components=2, perplexity=30). The max peak height per region was used to color each sequence.

### Cross-species cross-prediction correlations

We quantified group correspondence across species by correlating model predictions on matched, highly specific region sets and visualized per-cell-type correlations across species pairs. For each species AnnData, we selected the top-k (=500) most specific regions per cell type using *crested.pp.sort_and_filter_regions_on_specificity(method=“proportion”)*. For each species’ region set, we computed predictions with each species model (*crested.tl.predict*) to obtain cross-prediction tensors keyed by source species and cell type. For every pair of species (Mouse–Macaque, Mouse–Human, Macaque–Human), we formed class-by-class correlation matrices by comparing per-region prediction vectors between the two models, using Spearman correlation on log1p-transformed values; we also computed a “combined” matrix by concatenating both directional comparisons. We extracted the diagonal (same-name cell types) to summarize per-cell-type cross-species agreement.

### TF motif analysis

We derived cell type-specific sequence patterns for all species with their corresponding CREsted models using the contribution scores of the top 500 most specific regions per class. We selected these with *crested.pp.sort_and_filter_regions_on_specificity()* on the averaged peak heights and model predictions. For each region, we computed nucleotide-level contribution scores using CREsted’s *contribution_scores_specific()* function, method=’integrated_grad’. We ran *tfmodisco-lite*^50^ via *crested.tl.modisco.tfmodisco()* to discover recurrent patterns per group, and matched them to known motifs using the TOMTOM option (motifs from *crested.get_motif.db()*). Discovered patterns were post-processed and merged across groups using *crested.tl.modisco.process_patterns()*.

We matched motifs to TF candidates using the matched scRNA-seq data. We obtain pattern-to-motif matches from the motif database using *crested.tl.modisco.find_pattern_matches(q_val_thr=0.05)* and mapped motifs to TFs with *crested.tl.modisco.create_pattern_tf_dict()* using the human-oriented annotation columns. Finally, we constructed a TF x cell-type importance matrix with *crested.tl.modisco.create_tf_ct_matrix(min_tf_gex=0.95, importance_threshold=3, filter_correlation=True, zscore_threshold=1.5, correlation_threshold=0.49).* The resulting matrix summarizes, per cell type, TF candidates supported jointly by motif evidence and expression concordance.

### Evaluating *in-vivo* tested enhancers *in silico*

We scored 10 *in vivo* validated enhancers with the CREsted models. As the enhancers were variable in length, we zero-padded the flanks to obtain a 2,114 bp input sequence for the models. For the *TAC3* enhancer, we investigated its specificity by identifying all motifs within the sequence from the models’ contribution scores. To identify the motifs that define the enhancer’s Gluta-Dorsal-11&12 specificity, we destroyed each motif randomly by replacing it with a random sequence (n=20) and assessed that effect on the average delta prediction specificity of the mouse, human and macaque CREsted model. Specificity was defined as the proportion of the prediction on Gluta-Dorsal-11&12 and all the Gluta-Dorsal groups.

## Supplementary Tables

**Table S1. Taxonomy information**

**Table S2. Quality control metrics for consensus clusters**

**Table S3. Sample specimen information**

**Table S4. Macaque MERFISH gene panel**

**Table S5. Marker scores**

Per-species and conserved logistic regression scores markers for each Group, generated by scanpy’s rank_genes_groups function. The top 500 markers per Group are reported.

**Table S6. Mapping marker lists**

Marker lists used for correlation mapping across cell types in Figure 3D.

**Figure S1.**
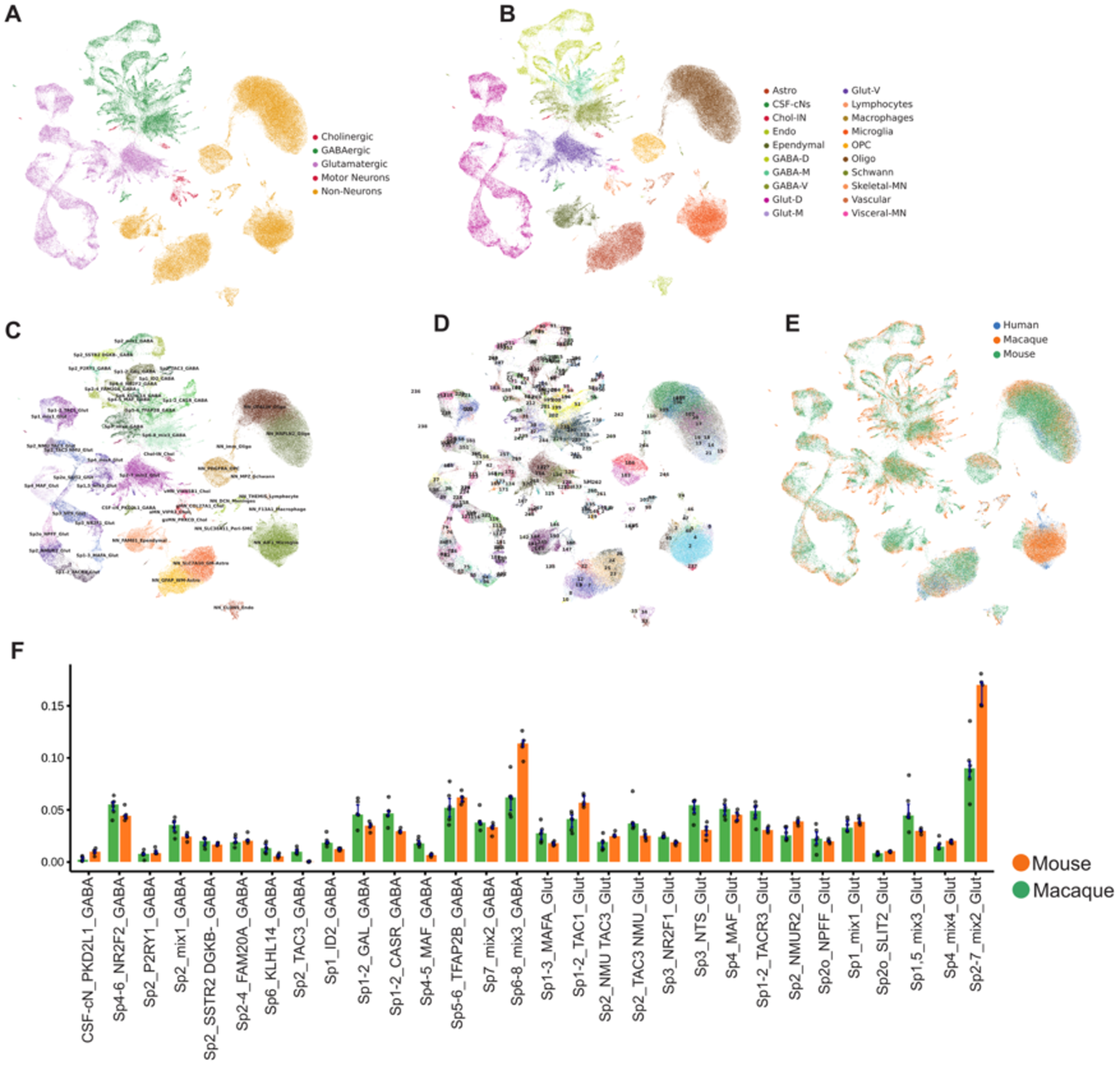
Additional UMAPs and neuron abundances, related to Figure 1. **(A-E)** UMAP projection of scVI cross-species latent space, colored by (A) Class, (B) Subclass, (C) Group, (D) consensus cluster, and (E) species. **(F)** Barplot of each Group’s proportion of total neurons in the dataset, split by species.

**Figure S2.**
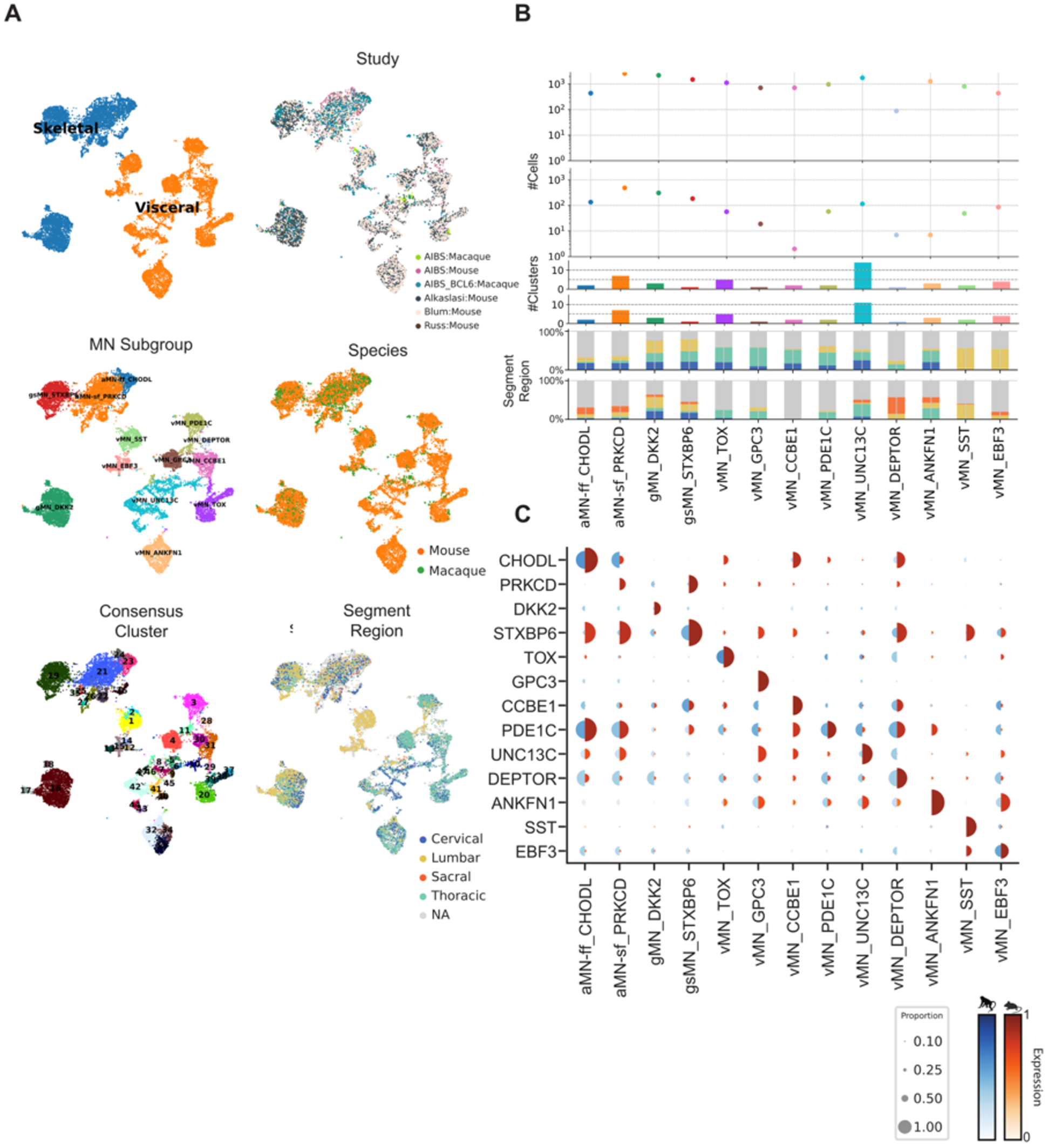
Motor neuron taxonomy, related to Figure 1. **(A)** UMAP projections of macaque and mouse motor neurons integrated using scVI, colored by taxonomy levels or metadata. **(B)** Atlas metadata of motor neuron taxonomy. Barplots represent the number of clusters within each MN Subgroup for each species. Dot plots represent the number of cells supporting each MN Subgroup from each species. **(C)** Pie dot-plot of the species-specific expression for selected conserved marker genes per Group term, where color palettes represent expression in each species, and dot diameter represents the proportion of cells in each Group with non-zero counts for that gene.

**Figure S3.**
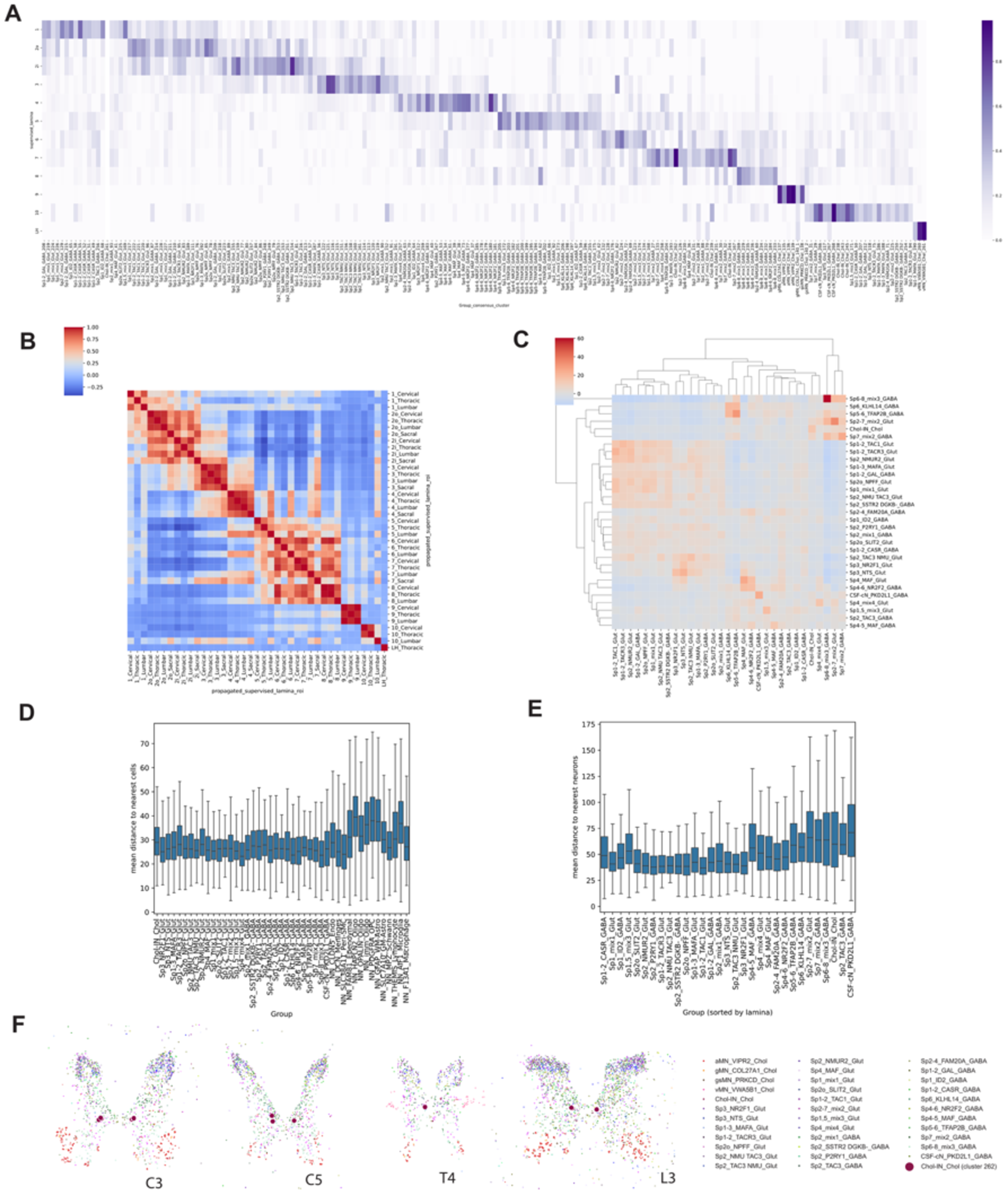
Additional macaque spatial transcriptomic analyses, related to Figure 2. **(A)** Heatmap showing the laminar distribution of consensus clusters in cervical, thoracic and lumbar segments. Values are cell counts normalized by total cells in each laminae, then across laminae for each Group. **(B)** Heatmap showing the Pearson correlation between the Group compositions of supervised laminae regions, for each segment. **(C)** Clustermap showing the Pearson correlation between the laminar distributions of each Group. **(D)** Boxplot showing the mean distance from each neuron to its nearest neighbors by Group. **(E)** Boxplot showing the mean distance from each cell (neurons and non-neurons) to its nearest neighbors by Group. **(F)** Plot showing locations of cells in spinal cord segments, colored by group with PITX2+ cholinergic INs (consensus cluster 262) enlarged to highlight their consistent location.

**Figure S4.**
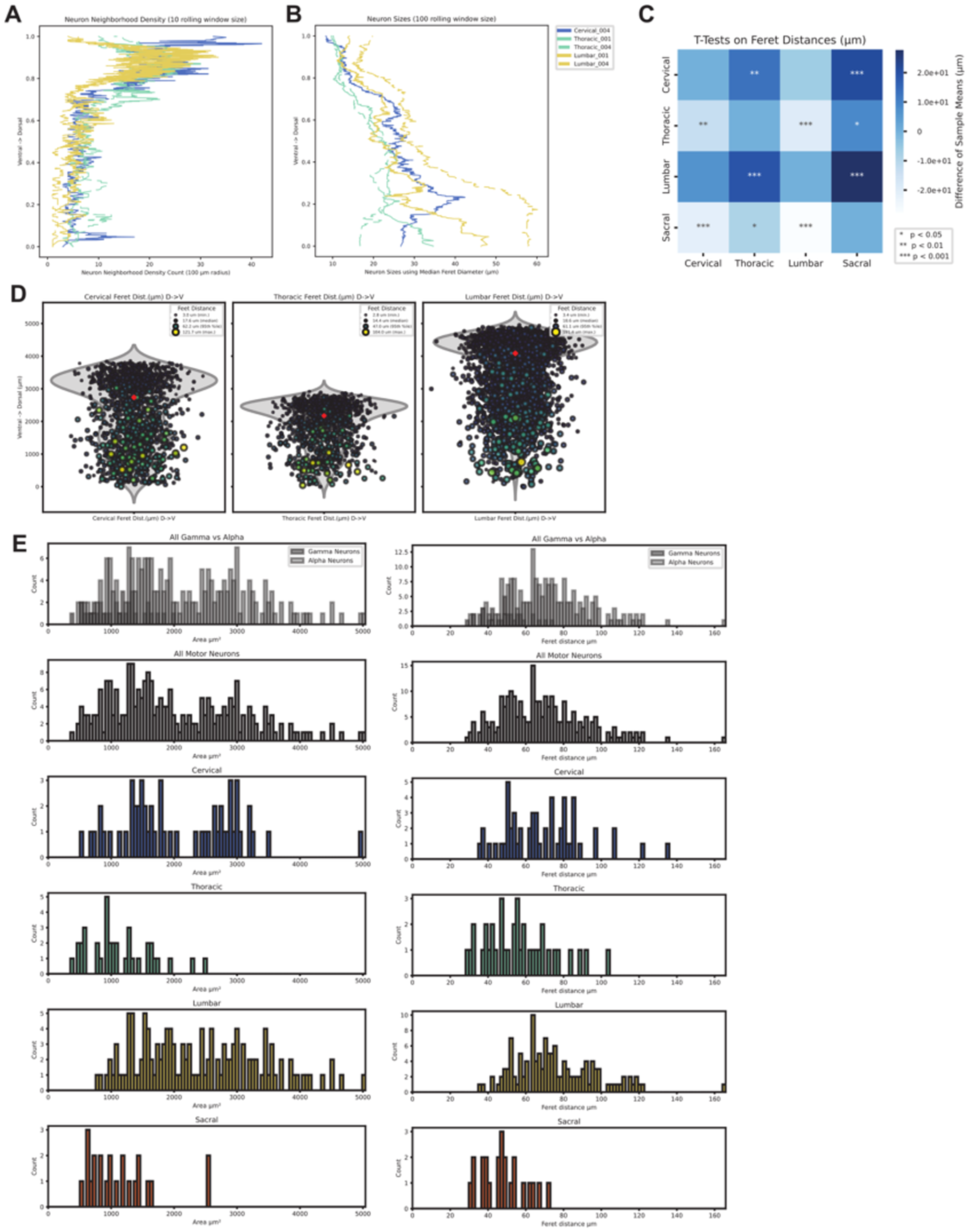
Additional macaque neuron size quantifications, related to Figure 2. **(A)** Lineplot showing the relationship between dorsal-ventral position and neuron density. **(B)** Lineplot showing the relationship between dorsal-ventral position and cell size. **(C)** Heatmap colored by pairwise t-tests across pairs for motor neuron Feret distances between spinal cord regions. **(D)** Stripplots with violin density plots showing the distribution of cells, with point size representing Feret distance. Mean values are represented by a red dot. **(E)** Histograms of motor neuron size per section (cervical, thoracic, lumbar, and sacral).

**Figure S5.**
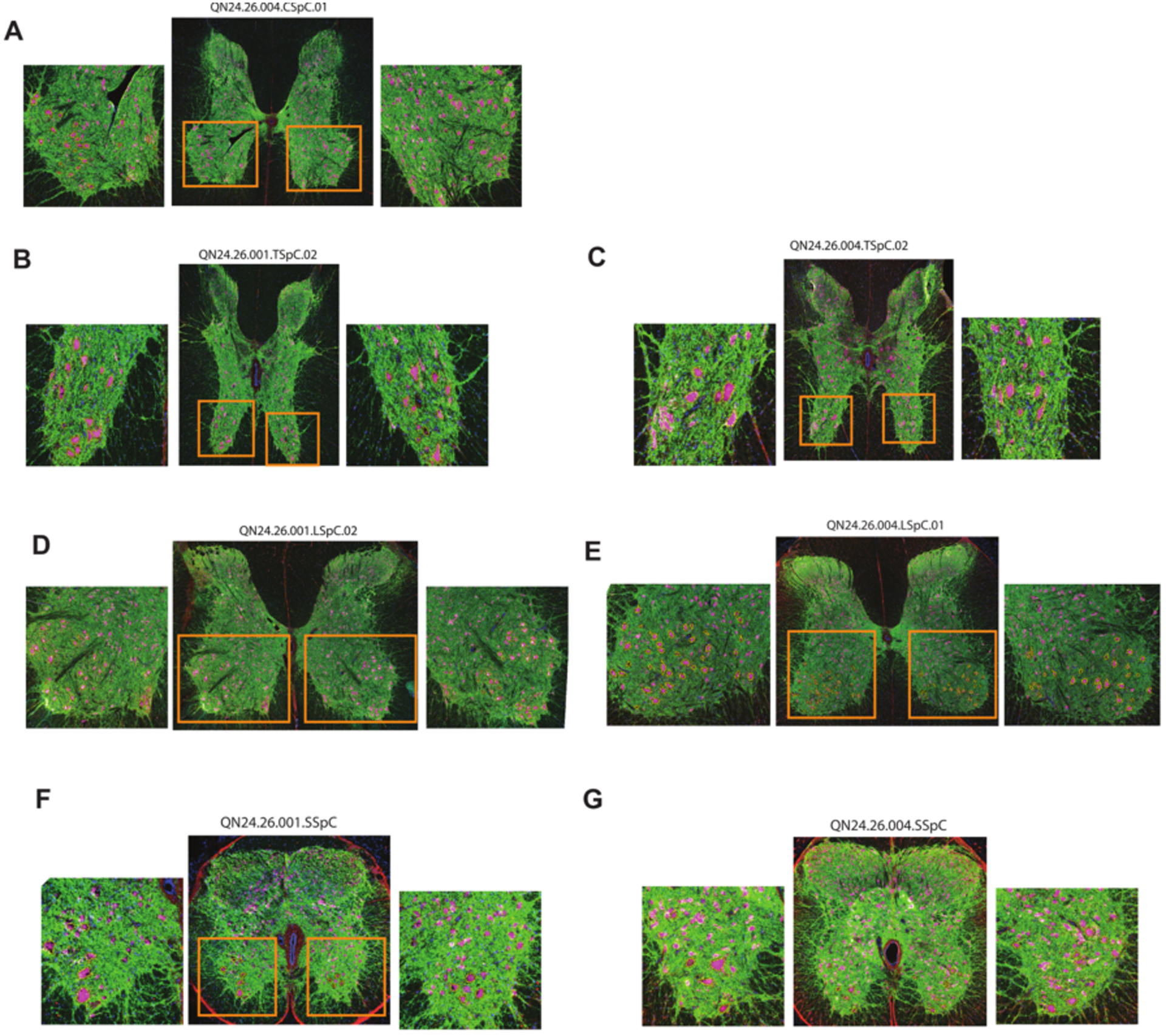
Immunofluorescence images, related to Figure 2. **(A-G)** Immunofluorescence images of macaque spinal sord sections tandem to those used for MERFISH spatial transcriptomics, zoom images on the ventral horn. Pseudocoloring is as follows: GFAP is green, NeuN is magenta, VGlut is red.

**Figure S6.**
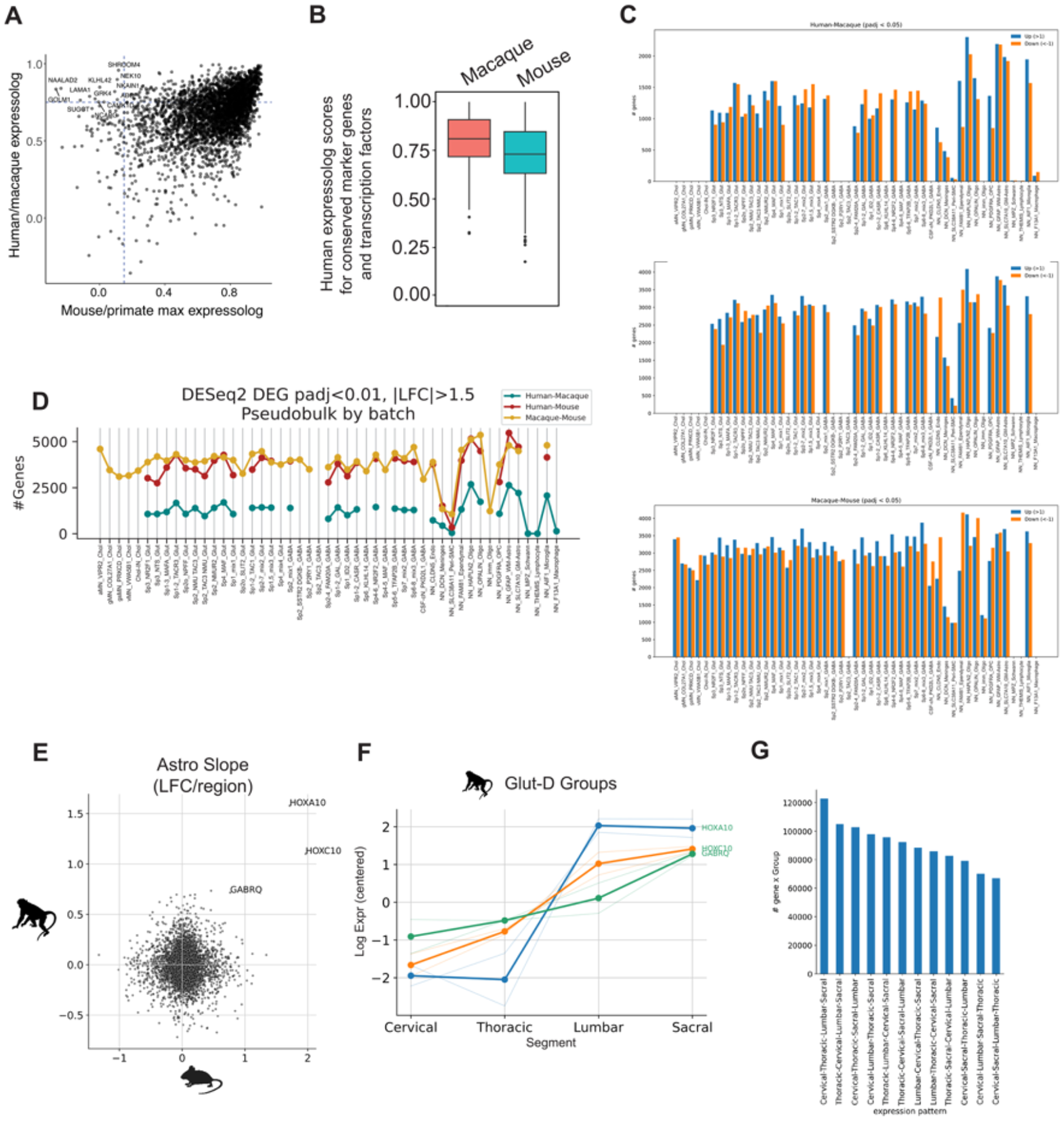
Additional cross-species transcriptome analyses, related to Figure 3. **(A)** Scatterplot showing the relationship between gene expression pattern correlation (expressolog) scores between species. Genes are labeled that were conserved (expressolog > 0.75) between human and macaque and divergent between mouse and both primate species (max expressolog < 0.2). **(B)** Boxplot showing the distribution of expressolog values relative to human for conserved marker genes and TFs showing expected higher conservation between human and macaque. **(C)** Barplot showing numbers of differentially expressed genes between pairs of species for each Group, with bar colors representing differentially expressed up vs down. **(D)** Lineplot showing numbers of differentially expressed genes between pairs of species for each Group, colored by species pairs. **(E)** Scatterplot of mean anterior-posterior gene expression slopes per gene (in log fold change per region) across cervical, thoracic, lumbar and sacral regions. Labeled genes have resampling p<0.1 for more than 50% of Groups in Subclass. **(F)** Lineplot showing the expression values of the significant genes from (E) (where lighter lines represent individual Astro Groups) **(G)** Barplot showing numbers of genes with expression means ordered by each possible region ordering showing the most common order is the rostro-caudal order (inverse orderings are merged with those shown, e.g. cervical, thoracic, lumbar, sacral versus sacral, lumbar, thoracic, cervical).

**Figure S7.**
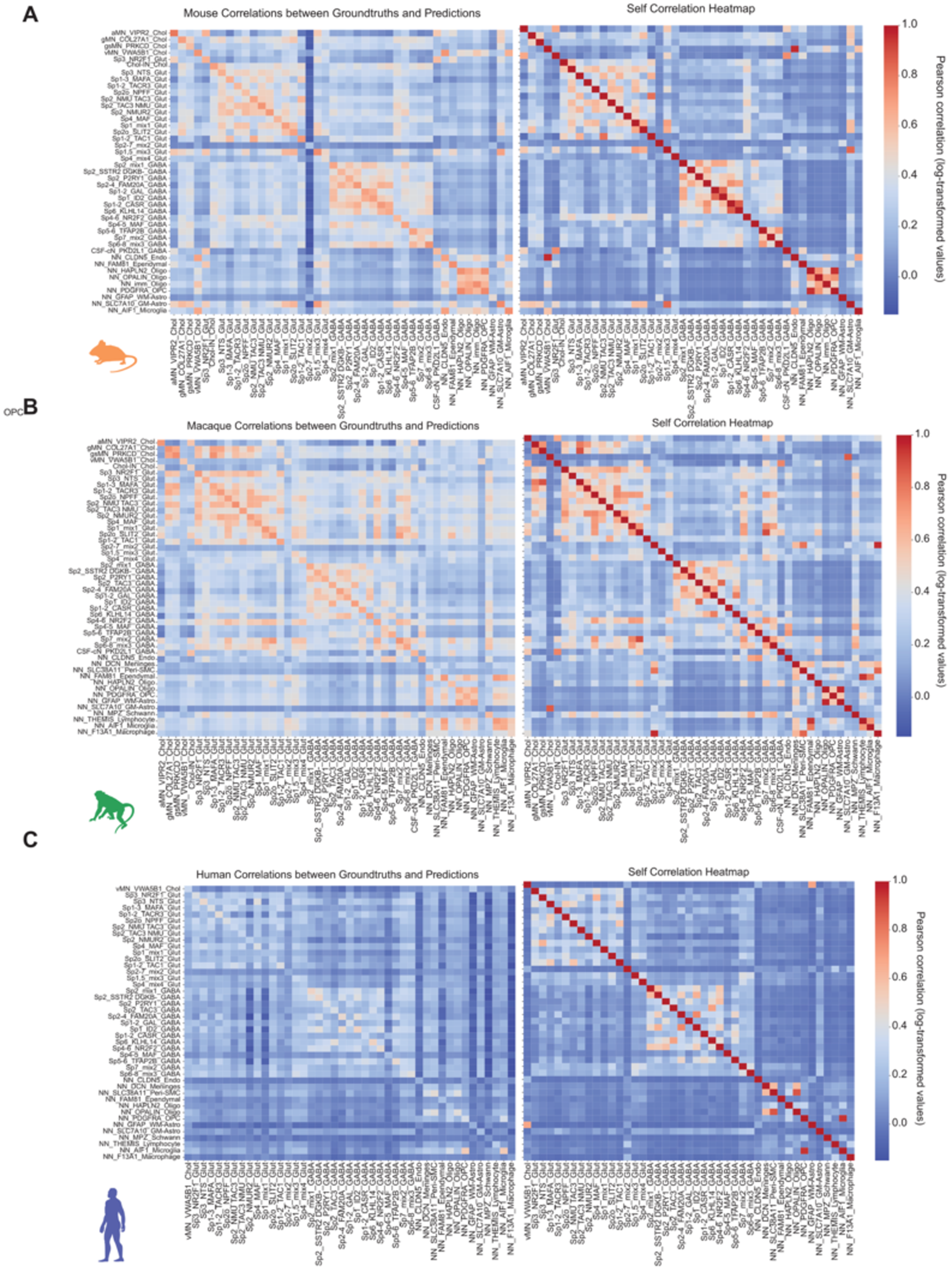
CREsted model prediction validation, related to Figure 5. **(A-C)** Heatmap colored by Pearson correlations of correlations between ground truth and CREsted-predicted (left) and ground truth (right) mean normalized log-transformed ATAC values from (A) mouse, (B) macaque, or (C) human single cell multiome data.

**Figure S8.**
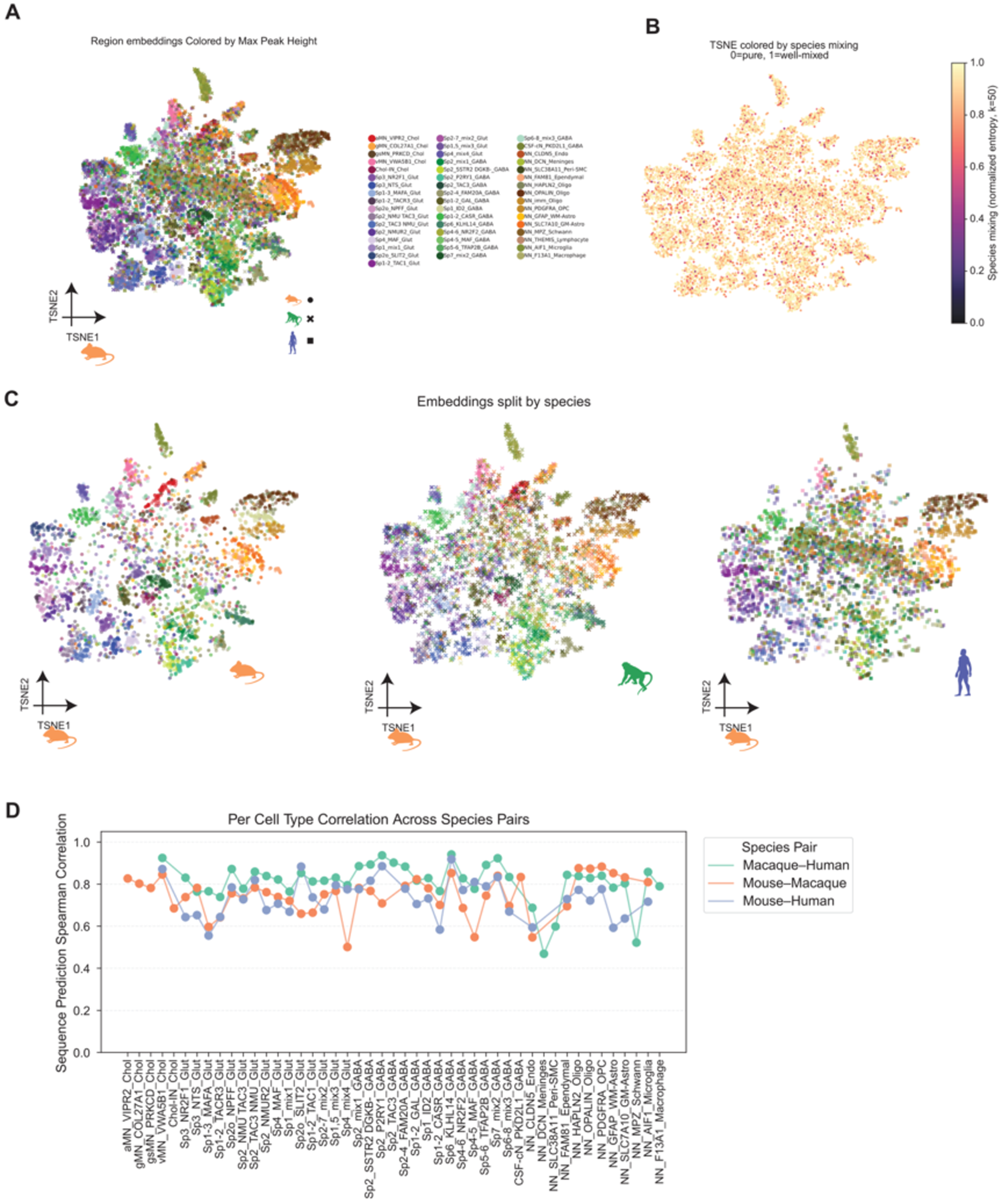
Mouse CREsted model predictions of peaks across species, related to Figure 5. **(A)** UMAP of DeepSpinalCord_mouse latent embeddings for the top 100 Group-specific cCREs from each species colored by the Group to which each peak is specific. **(B)** UMAP from panel (A) colored by species entropy values among k-nearest neighbors where 1 represents perfect mixing between species and 0 represents perfect species segregation. This shows that Groups lack species-specific features that are represented by the DeepSpinalCord_mouse model. **(C)** UMAP from panel (A) colored by the Group to which each peak is specific and split by each species. **(D)** Line plot showing the per Group Spearman correlation of peak openness predicted for each species pair.

## REFERENCES

1. Lemon, R.N., and Griffiths, J. (2005). Comparing the function of the corticospinal system in different species: organizational differences for motor specialization? Muscle Nerve 32, 261–279.

2. Toossi, A., Bergin, B., Marefatallah, M., Parhizi, B., Tyreman, N., Everaert, D.G., Rezaei, S., Seres, P., Gatenby, J.C., Perlmutter, S.I., et al. (2021). Comparative neuroanatomy of the lumbosacral spinal cord of the rat, cat, pig, monkey, and human. Sci. Rep. 11, 1955.

3. Apkarian, A.V., and Hodge, C.J. (1989). Primate spinothalamic pathways: II. The cells of origin of the dorsolateral and ventral spinothalamic pathways. J. Comp. Neurol. 288, 474–492.

4. Zhang, E.T., and Craig, A.D. (1997). Morphology and distribution of spinothalamic lamina I neurons in the monkey. J. Neurosci. 17, 3274–3284.

5. Arokiaraj, C.M., Leone, M.J., Kleyman, M., Chamessian, A., Noh, M.-C., Phan, B.N., Lopes, B.C., Corrigan, K.A., Cherupally, V.K., Yeramosu, D., et al. (2024). Spatial, transcriptomic, and epigenomic analyses link dorsal horn neurons to chronic pain genetic predisposition. Cell Rep. 43, 114876.

6. Yadav, A., Matson, K.J.E., Li, L., Hua, I., Petrescu, J., Kang, K., Alkaslasi, M.R., Lee, D.I., Hasan, S., Galuta, A., et al. (2023). A cellular taxonomy of the adult human spinal cord. Neuron 111, 328–344.e7.

7. Siletti, K., Hodge, R., Mossi Albiach, A., Lee, K.W., Ding, S.-L., Hu, L., Lönnerberg, P., Bakken, T., Casper, T., Clark, M., et al. (2023). Transcriptomic diversity of cell types across the adult human brain. Science 382, eadd7046.

8. Yao, Z., van Velthoven, C.T.J., Kunst, M., Zhang, M., McMillen, D., Lee, C., Jung, W., Goldy, J., Abdelhak, A., Aitken, M., et al. (2023). A high-resolution transcriptomic and spatial atlas of cell types in the whole mouse brain. Nature 624, 317–332.

9. Jorstad, N.L., Song, J.H.T., Exposito-Alonso, D., Suresh, H., Castro-Pacheco, N., Krienen, F.M., Yanny, A.M., Close, J., Gelfand, E., Long, B., et al. (2023). Comparative transcriptomics reveals human-specific cortical features. Science 382, eade9516.

10. Ma, S., Skarica, M., Li, Q., Xu, C., Risgaard, R.D., Tebbenkamp, A.T.N., Mato-Blanco, X., Kovner, R., Krsnik, Ž., de Martin, X., et al. (2022). Molecular and cellular evolution of the primate dorsolateral prefrontal cortex. Science, eabo7257.

11. Hodge, R.D., Bakken, T.E., Miller, J.A., Smith, K.A., Barkan, E.R., Graybuck, L.T., Close, J.L., Long, B., Johansen, N., Penn, O., et al. (2019). Conserved cell types with divergent features in human versus mouse cortex. Nature 573, 61–68.

12. Russ, D.E., Cross, R.B.P., Li, L., Koch, S.C., Matson, K.J.E., Yadav, A., Alkaslasi, M.R., Lee, D.I., Le Pichon, C.E., Menon, V., et al. (2021). A harmonized atlas of mouse spinal cord cell types and their spatial organization. Nat. Commun. 12, 5722.

13. Alkaslasi, M.R., Piccus, Z.E., Hareendran, S., Silberberg, H., Chen, L., Zhang, Y., Petros, T.J., and Le Pichon, C.E. (2021). Single nucleus RNA-sequencing defines unexpected diversity of cholinergic neuron types in the adult mouse spinal cord. Nat. Commun. 12, 2471.

14. Blum, J.A., Klemm, S., Shadrach, J.L., Guttenplan, K.A., Nakayama, L., Kathiria, A., Hoang, P.T., Gautier, O., Kaltschmidt, J.A., Greenleaf, W.J., et al. (2021). Single-cell transcriptomic analysis of the adult mouse spinal cord reveals molecular diversity of autonomic and skeletal motor neurons. Nat. Neurosci. 24, 572–583.

15. Gautier, O., Blum, J.A., Maksymetz, J., Chen, D., Schweingruber, C., Mei, I., Hermann, A., Hackos, D.H., Hedlund, E., Ravits, J., et al. (2023). Challenges of profiling motor neuron transcriptomes from human spinal cord. Neuron 111, 3739–3741.

16. Gautier, O., Blum, J.A., Nguyen, T.P., Klemm, S., Yamakawa, M., Sinnott-Armstrong, N., Zeng, Y., Davis, C.-H.O., Bombosch, J., Nakayama, L., et al. (2025). An emergent disease-associated motor neuron state precedes cell death in a mouse model of ALS. bioRxivorg, 2025.08.21.671404. 10.1101/2025.08.21.671404.

17. Johansen, N.J., Fu, Y., Schmitz, M., Dubuc, A., Kempynck, N., Wirthlin, M., Garcia, A.D., Hewitt, M., Turner, M.A., Seeman, S.C., et al. (2025). Cross-species consensus atlas of the primate basal ganglia. bioRxivorg, 2025.12.15.694496. 10.64898/2025.12.15.694496.

18. Corrigan, E.K., DeBerardine, M., Poddar, A., Turrero García, M., de la O, S., He, S., Sen, H., Duhne, M., Lindberg, S., Song, M., et al. (2025). Conservation and alteration of mammalian striatal interneurons. Nature 647, 187–193.

19. Bakken, T.E., van Velthoven, C.T., Menon, V., Hodge, R.D., Yao, Z., Nguyen, T.N., Graybuck, L.T., Horwitz, G.D., Bertagnolli, D., Goldy, J., et al. (2021). Single-cell and single-nucleus RNA-seq uncovers shared and distinct axes of variation in dorsal LGN neurons in mice, non-human primates, and humans. Elife 10. 10.7554/eLife.64875.

20. Affinati, A.H., Sabatini, P.V., True, C., Tomlinson, A.J., Kirigiti, M., Lindsley, S.R., Li, C., Olson, D.P., Kievit, P., Myers, M.G., et al. (2021). Cross-species analysis defines the conservation of anatomically segregated VMH neuron populations. Elife 10. 10.7554/eLife.69065.

21. Hahn, J., Monavarfeshani, A., Qiao, M., Kao, A.H., Kölsch, Y., Kumar, A., Kunze, V.P., Rasys, A.M., Richardson, R., Wekselblatt, J.B., et al. (2023). Evolution of neuronal cell classes and types in the vertebrate retina. Nature 624, 415–424.

22. Sepp, M., Leiss, K., Murat, F., Okonechnikov, K., Joshi, P., Leushkin, E., Spänig, L., Mbengue, N., Schneider, C., Schmidt, J., et al. (2024). Cellular development and evolution of the mammalian cerebellum. Nature 625, 788–796.

23. Fu, Y., Johansen, N.J., Kempynck, N., Ding, W., Turner, M.A., Garcia, A.D., Schmitz, M.T., Close, J., Kapen, I., Hewitt, M., et al. (2025). Circuit specific specialization of human basal ganglia astrocytes. bioRxiv. 10.64898/2025.12.19.695583.

24. Landa-García, J.N., Palacios-Arellano, M. de la P., Morales, M.A., Aranda-Abreu, G.E., Rojas-Durán, F., Herrera-Covarrubias, D., Toledo-Cárdenas, M.R., Suárez-Medellín, J.M., Coria-Avila, G.A., Manzo, J., et al. (2024). The anatomy, histology, and function of the major pelvic ganglion. Animals (Basel) 14, 2570.

25. Baek, M., Menon, V., Jessell, T.M., Hantman, A.W., and Dasen, J.S. (2019). Molecular Logic of Spinocerebellar Tract Neuron Diversity and Connectivity. Neuroscience.

26. Zagoraiou, L., Akay, T., Martin, J.F., Brownstone, R.M., Jessell, T.M., and Miles, G.B. (2009). A cluster of cholinergic premotor interneurons modulates mouse locomotor activity. Neuron 64, 645–662.

27. Zhang, M., Eichhorn, S.W., Zingg, B., Yao, Z., Cotter, K., Zeng, H., Dong, H., and Zhuang, X. (2021). Spatially resolved cell atlas of the mouse primary motor cortex by MERFISH. Nature 598, 137–143.

28. Hardesty, I. (1902). Observations on the medulla spinalis of the elephant with some comparative studies of the intumescentia cervicalis and the neurones of the columna anterior. J. Comp. Neurol. 12, 125–182.

29. Manuel, M., Chardon, M., Tysseling, V., and Heckman, C.J. (2019). Scaling of motor output, from mouse to humans. Physiology (Bethesda) 34, 5–13.

30. Shneider, N.A., Brown, M.N., Smith, C.A., Pickel, J., and Alvarez, F.J. (2009). Gamma motor neurons express distinct genetic markers at birth and require muscle spindle-derived GDNF for postnatal survival. Neural Dev. 4, 42.

31. Friese, A., Kaltschmidt, J.A., Ladle, D.R., Sigrist, M., Jessell, T.M., and Arber, S. (2009). Gamma and alpha motor neurons distinguished by expression of transcription factor Err3. Proc. Natl. Acad. Sci. U. S. A. 106, 13588–13593.

32. Cullheim, S., and Ulfhake, B. (1979). Relations between cell body size, axon diameter and axon conduction velocity of triceps surae alpha montoneurons during the postnatal development in the cat. J. Comp. Neurol. 188, 679–686.

33. Jorstad, N.L., Close, J., Johansen, N., Yanny, A.M., Barkan, E.R., Travaglini, K.J., Bertagnolli, D., Campos, J., Casper, T., Crichton, K., et al. (2023). Transcriptomic cytoarchitecture reveals principles of human neocortex organization. Science 382, eadf6812.

34. Fang, Z., Liu, X., and Peltz, G. (2023). GSEApy: a comprehensive package for performing gene set enrichment analysis in Python. Bioinformatics 39, btac757.

35. Kuderna, L.F.K., Ulirsch, J.C., Rashid, S., Ameen, M., Sundaram, L., Hickey, G., Cox, A.J., Gao, H., Kumar, A., Aguet, F., et al. (2024). Identification of constrained sequence elements across 239 primate genomes. Nature 625, 735–742.

36. Suresh, H., Crow, M., Jorstad, N., Hodge, R., Lein, E., Dobin, A., Bakken, T., and Gillis, J. (2023). Comparative single-cell transcriptomic analysis of primate brains highlights human-specific regulatory evolution. Nat. Ecol. Evol. 7, 1930–1943.

37. Kempynck, N., De Winter, S., Blaauw, C.H., Konstantakos, V., Dieltiens, S., Eksi, E.Ç., Bercier, V., Taskiran, I.I., Hulselmans, G., Spanier, K., et al. (2025). CREsted: modeling genomic and synthetic cell type-specific enhancers across tissues and species. bioRxiv, 2025.04.02.646812. 10.1101/2025.04.02.646812.

38. Kussick, E., Johansen, N., Taskin, N., Chowdhury, A., Quinlan, M.A., Fraser, A., Clark, A.G., Wynalda, B., Martinez, R., Groce, E.L., et al. (2025). Enhancer AAVs for targeting spinal motor neurons and descending motor pathways in rodents and macaque. Cell Rep. 44, 115730.

39. Chen, S., Gao, X.-F., Zhou, Y., Liu, B.-L., Liu, X.-Y., Zhang, Y., Barry, D.M., Liu, K., Jiao, Y., Bardoni, R., et al. (2020). A spinal neural circuitry for converting touch to itch sensation. Nat. Commun. 11, 5074.

40. Schmitz, M.T., Ding, J.W., Nolbrant, S., McMullen, R., Kim, C.N., Pavlovic, B.J., Nowakowski, T.J., Bakken, T.E., Ye, C.J., and Pollen, A.A. (2025). ANTIPODE provides a global view of cell type homology and transcriptomic divergence in the developing mammalian brain. bioRxivorg. 10.1101/2025.10.18.683238.

41. Chong, A., Song, H.-C., Byun, B.-H., Hong, S.-P., Min, J.-J., Bom, H.-S., Ha, J.-M., and Lee, J.-K. (2013). Changes in (18)f-fluorodeoxyglucose uptake in the spinal cord in a healthy population on serial positron emission tomography/computed tomography. Chonnam Med. J. 49, 38–42.

42. Tao, S., Smith, K.A., Ng, L., Li, X., Huang, Y., Mufti, S., Ghosh, S.S., Martone, M.E., White, O., and Zhang, G. (2025). Tissue-to-Bytes: A Catalytic Digital Twin Platform for Consortium-Scale Integration of Single-Cell Omics Data Across the BRAIN Initiative Cell Atlas Network. Preprint.

43. Ding, S.-L., and Lein, E.S. Towards human and non-human primate common coordinate frameworks using unified structural ontologies. Preprint.

44. Crow, M., Paul, A., Ballouz, S., Huang, Z.J., and Gillis, J. (2018). Characterizing the replicability of cell types defined by single cell RNA-sequencing data using MetaNeighbor. Nat. Commun. 9, 884.

45. Hewitt, M.N., Turner, M.A., Johansen, N., McMillen, D.A., Dan, S., DeBerardine, M., Ruiz, A., Huang, M., Quon, J., Fu, Y., et al. (2025). A cross-species spatial transcriptomic atlas of the human and non-human primate basal ganglia. Neuroscience.

46. Zhang, K., Zemke, N.R., Armand, E.J., and Ren, B. (2024). A fast, scalable and versatile tool for analysis of single-cell omics data. Nat. Methods 21, 217–227.

47. Yanai, I., Benjamin, H., Shmoish, M., Chalifa-Caspi, V., Shklar, M., Ophir, R., Bar-Even, A., Horn-Saban, S., Safran, M., Domany, E., et al. (2005). Genome-wide midrange transcription profiles reveal expression level relationships in human tissue specification. Bioinformatics 21, 650–659.

48. Paten, B., Earl, D., Nguyen, N., Diekhans, M., Zerbino, D., and Haussler, D. (2011). Cactus: Algorithms for genome multiple sequence alignment. Genome Res. 21, 1512–1528.

49. Zhang, X., Kaplow, I.M., Wirthlin, M., Park, T.Y., and Pfenning, A.R. (2020). HALPER facilitates the identification of regulatory element orthologs across species. Bioinformatics 36, 4339–4340.

50. Wang, Tseng, Ramalingam, Schreiber, and co-authors (2026). Decoding predictive motif lexicons and syntax from deep learning models of transcription factor binding profiles (in preparation).

